# Ligand-Induced Structural Dynamics Drive Allosteric Regulation of Translation Initiation Factor eIF4E

**DOI:** 10.64898/2025.12.23.696204

**Authors:** Agnese Roscioni, Vincenzo Maria D’Amore, Jesmina Rexha, Greta R. Bianchini, Anna La Teana, Daniele Di Marino, Alice Romagnoli, Francesco Saverio Di Leva

## Abstract

The eukaryotic initiation factor 4E (eIF4E) governs cap-dependent translation, and its dysregulation contributes both to cancer and neurological disorders, making it an attractive target for therapeutic intervention. Among the few small-molecule inhibitors developed, 4EGI-1 and its analogue i4EG-BiP have shown promise in cellular and preclinical models. These compounds disrupt eIF4E’s interaction with its partner eIF4G while enhancing binding to its negative regulators, the 4E-binding proteins (4E-BPs). Despite their chemical similarity and overlapping functional effects, structural data suggest the two ligands engage different regions of eIF4E, lateral for 4EGI-1 and frontal for i4EG-BiP, raising questions about the basis of their shared activity.

Here, we integrate molecular simulations, site-directed mutagenesis, and fluorescence binding assays to elucidate the mechanism of action of both ligands. Funnel metadynamics free-energy calculations reveal that the frontal binding mode is thermodynamically preferred for both compounds. Furthermore, we demonstrate that the conformational rearrangement induced by frontal binding selectively promotes 4E-BP1 over eIF4G association, explaining their common allosteric regulatory effect.

These findings reconcile divergent structural observations and highlight how ligand-induced dynamics can be exploited to reprogram eIF4E interactions, offering a framework for next-generation therapeutics targeting dysregulated translation.

## INTRODUCTION

Translation initiation is a fundamental cellular process intricately regulated by a network of molecular factors, among which the Eukaryotic Initiation Factor 4E (eIF4E) plays a pivotal role. eIF4E is a 25 kDa horseshoe-shaped protein composed of eight antiparallel β-strands and three elongated α-helices forming distinct ventral, dorsal, frontal and lateral surfaces (Figure 1A)^1^. Together with the DEAD-box helicase eIF4A and the scaffold protein eIF4G, eIF4E assembles into the heterotrimeric eIF4F complex, a crucial regulator of translation initiation^2–6^. Acting as a molecular bridge, eIF4F recruits the 43S Pre-Initiation Complex (43S PIC) to the 5’ end of the mRNA. In this assembly, eIF4E binds the 7-methylguanosine (m^7^GTP) cap, eIF4G connects to the 43S PIC via eIF3, and eIF4A unwinds secondary structures in the 5’ untranslated region (UTR) to enable efficient ribosomal scanning^2,3^.

**Figure 1.**
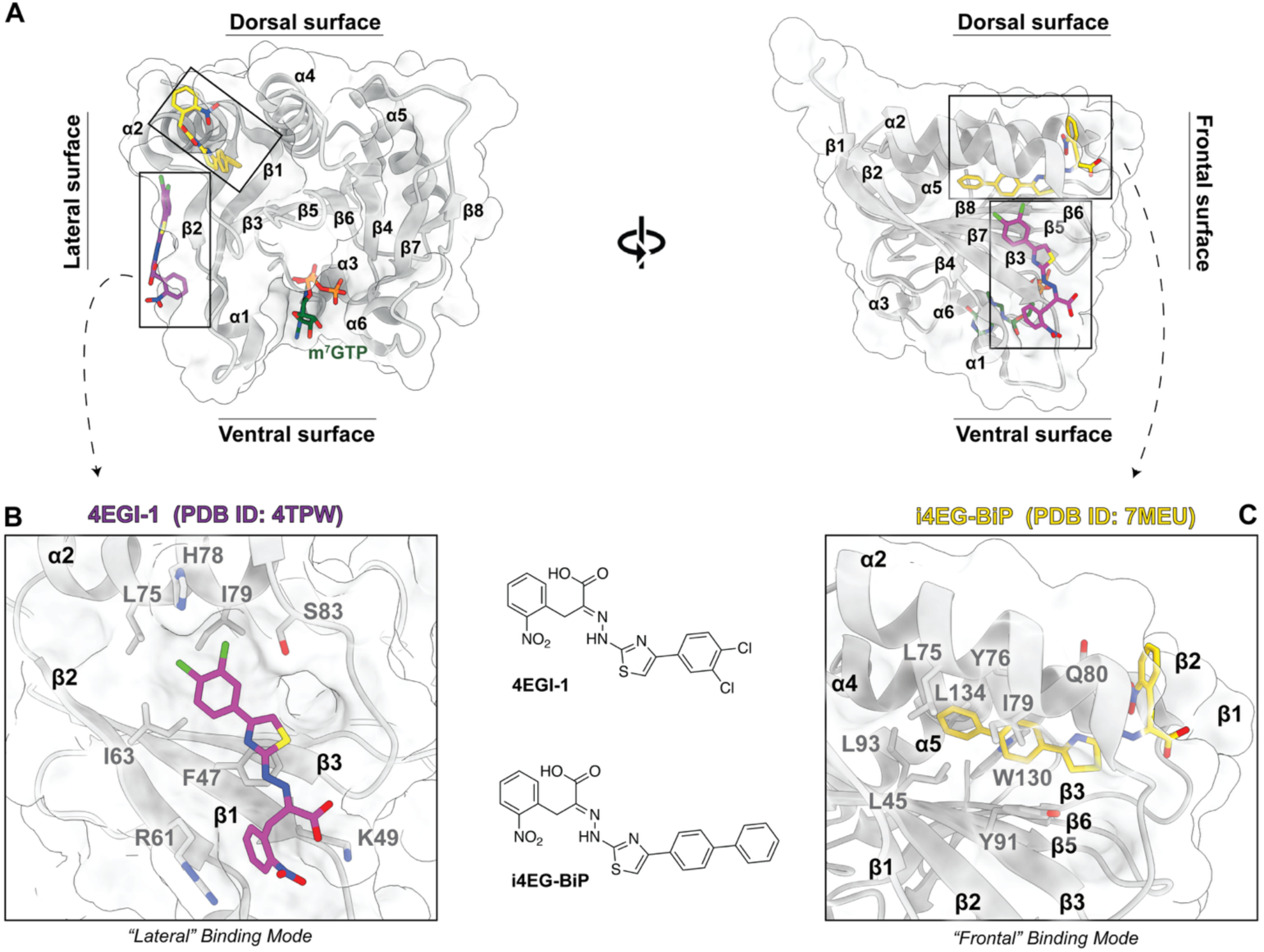
4EGI-1 and i4EG-BiP experimental binding modes. A) Three-dimensional structure of eIF4E (PDB ID: 4TPW) shown from two orientations (rotated by 90°) to visualize both the lateral (left) and frontal (right) surfaces. Experimental binding sites of 4EGI-1 (magenta sticks), i4EG-BiP (yellow sticks), and m^7^GTP (green sticks) are highlighted. B-C) Close-up views of the lateral (B) and frontal (C) binding modes adopted by 4EGI-1 (PDB ID: 4TPW) and i4EG-BiP (PDB ID: 7MEU), respectively. Key residues involved in ligand binding are shown as sticks. The chemical structures of the two compounds are also provided. Non-polar hydrogens are omitted for clarity.

Among eIF4F components, eIF4E is the least abundant, imposing a quantitative constraint on complex formation^7,8^. Its expression and activity are tightly regulated through transcriptional, post-transcriptional, and post-translational mechanisms, including its interaction with 4E-binding proteins (4E-BPs)^1^. These are established negative modulators of translation initiation, as they compete with eIF4G for binding to eIF4E, thereby preventing the formation of the eIF4F complex^1,9–11^. Aberrant regulation of eIF4E is a hallmark of numerous tumor types, including breast cancer, non-Hodgkin lymphoma, and head and neck cancer^12–17^. Notably, targeting of signaling pathways converging on eIF4E has demonstrated synergistic effects with current immunotherapies, contributing to the restoration of immune surveillance in prostate cancer^18,19^. Moreover, it has been implicated in neurological disorders, such as Autism Spectrum Disorders (ASD) and Fragile X Syndrome (FXS)^1,20,21^. As a result, eIF4E has emerged as a promising therapeutic target ^1,22–27^, and multiple strategies have been developed to inhibit its function, including cap analogs, peptides, RNA-based approaches and small molecules^3,28–38^. Among these, 4EGI-1 and its biphenyl derivative i4EG-BiP have shown promising preclinical efficacy across various cancer cell lines^39–41^, and in ASD and FXS mouse models^21,42^. Both compounds inhibit the eIF4E/eIF4G interaction with comparable potency^40,41,43^, while enhancing the interaction between eIF4E and 4E-BPs^39^. The molecular basis of their interaction with eIF4E was first revealed upon the release of two crystallographic complexes. These structures show 4EGI-1 bound at the lateral surface of eIF4E (PDB ID: 4TPW^40^), adjacent to helix α1, and i4EG-BiP bound to a frontal pocket between helices α2 and α4 (PDB ID: 7MEU^41^) corresponding to the “site 2” recently identified by Clarke and co-workers^32^. We refer to these as the “lateral” and “frontal” binding modes, respectively (Figure 1A-C). X-ray complexes show that the two compounds bind to different sites on eIF4E, despite having similar chemical and pharmacological features. This stands in contrast to the nearly identical ability to disrupt eIF4G binding and promote the association with 4E-BPs. This convergence of activity would in fact suggest a common mechanism of action that is not readily explained by existing static structural models^9^. Furthermore, both ligand-bound structures exhibit a distinct conformational change in eIF4E, absent in the apo state and in complexes with eIF4G or 4E-BPs, pointing to a ligand-induced effect with potential regulatory implications^40,41^. While crystallographic^40,41^, biochemical^44^, and mutational^40^ data have provided valuable insights, they are insufficient to clarify how the two ligands can produce the same outcome occupying distinct sites. Moreover, a comparative analysis of the dynamic and energetic features of the two binding modes is still lacking.

In this study, we address these unresolved questions by combining computational and experimental approaches to elucidate, at molecular resolution, the interaction of 4EGI-1 and i4EG-BiP with eIF4E.

Through extensive molecular dynamics (MD) simulations and free energy calculations using funnel metadynamics (FM)^45,46^, we characterize the stability and dynamics of the two crystallographic binding poses. Our results demonstrate that the frontal binding mode is energetically and dynamically favored for both ligands, providing further insights into their inhibitory mechanism. We further clarify how binding of 4EGI-1 and i4EG-BiP modulates the mutually exclusive association of eIF4E with eIF4G and 4E-BPs, by analyzing ternary eIF4E–ligand complexes with these partners. These findings reconcile structural and functional data and uncover a shared mechanism of allosteric regulation, in which ligand binding impairs eIF4G recruitment while enhancing 4E-BPs association. The computational predictions are further supported by targeted mutagenesis experiments, reinforcing the proposed model of ligand-induced modulation. Overall, our study sheds light on the molecular basis of eIF4E inhibition and offers a solid framework for the rational design of next-generation modulators of this critical translation factor.

## RESULTS

### Molecular Simulations Support Common Binding Mode for 4EGI-1 and i4EG-BiP

As the first step in our investigation of the binding dynamics of 4EGI-1 and i4EG-BiP with eIF4E, we carried out all-atom MD simulations in explicit solvent starting from their respective crystallographic complexes (PDB IDs: 4TPW and 7MEU). The lateral pose assumed by 4EGI-1 (Figure 1B), is primarily stabilized by: (i) a salt bridge between the ligand’s carboxylate group and the K49 side chain; (ii) a cation-π interaction between the *o*-nitrophenyl group and the R61 guanidine group; and (iii) stacking interactions between the thiazole ring and the F47 side chain^40^. In the frontal pose assumed by i4EG-BiP (Figure 1C), the ligand’s biphenyl moiety is deeply buried within the cavity defined by the α2 and α4 helices, where it forms edge-to-face π-π interactions with Y76, Y91, and W130. The nitrophenyl, hydrazone, and carboxylic acid groups remain solvent-exposed, forming H-bonds with Q80. Additional hydrophobic contacts are observed between the ligand and residues L45, L75, I79, L93, and L134 (Fig 1C)^41^. To ensure statistical robustness, each pose was simulated through four independent replicas of 3 μs, yielding a total simulation time of 12 μs per system. This approach enabled a comprehensive examination of the ligand-protein interactions, fully accounting for full receptor flexibility and solvent effects.

Notable differences in the behavior of the two ligands emerge early when assessing the temporal stability of their initial binding poses. Specifically, 4EGI-1 exhibits high fluctuations, failing to maintain its binding mode and experiencing unbinding events across the four simulated replicas. In contrast, i4EG-BiP demonstrates significantly greater stability, consistently retaining its starting conformation throughout all trajectories. This qualitative distinction becomes even more evident when tracking the movement of the ligands’ center of mass (CoM) over the course of the simulations. As illustrated in Figure 2, the CoM of 4EGI-1 (magenta spheres, Figure 2A) is dispersed across the protein surface and frequently shifts into the bulk solvent. In stark contrast, the CoM of i4EG-BiP (yellow spheres, Figure 2C) remains tightly clustered within the initial binding site. A more quantitative evaluation of this phenomenon is provided by root mean square deviation (RMSD) analysis (Figure 2B and 2D), demonstrating higher fluctuations for 4EGI-1. These data are consistent with the reduced stability of its lateral binding mode compared to the frontal one adopted by i4EG-BiP.

**Figure 2.**
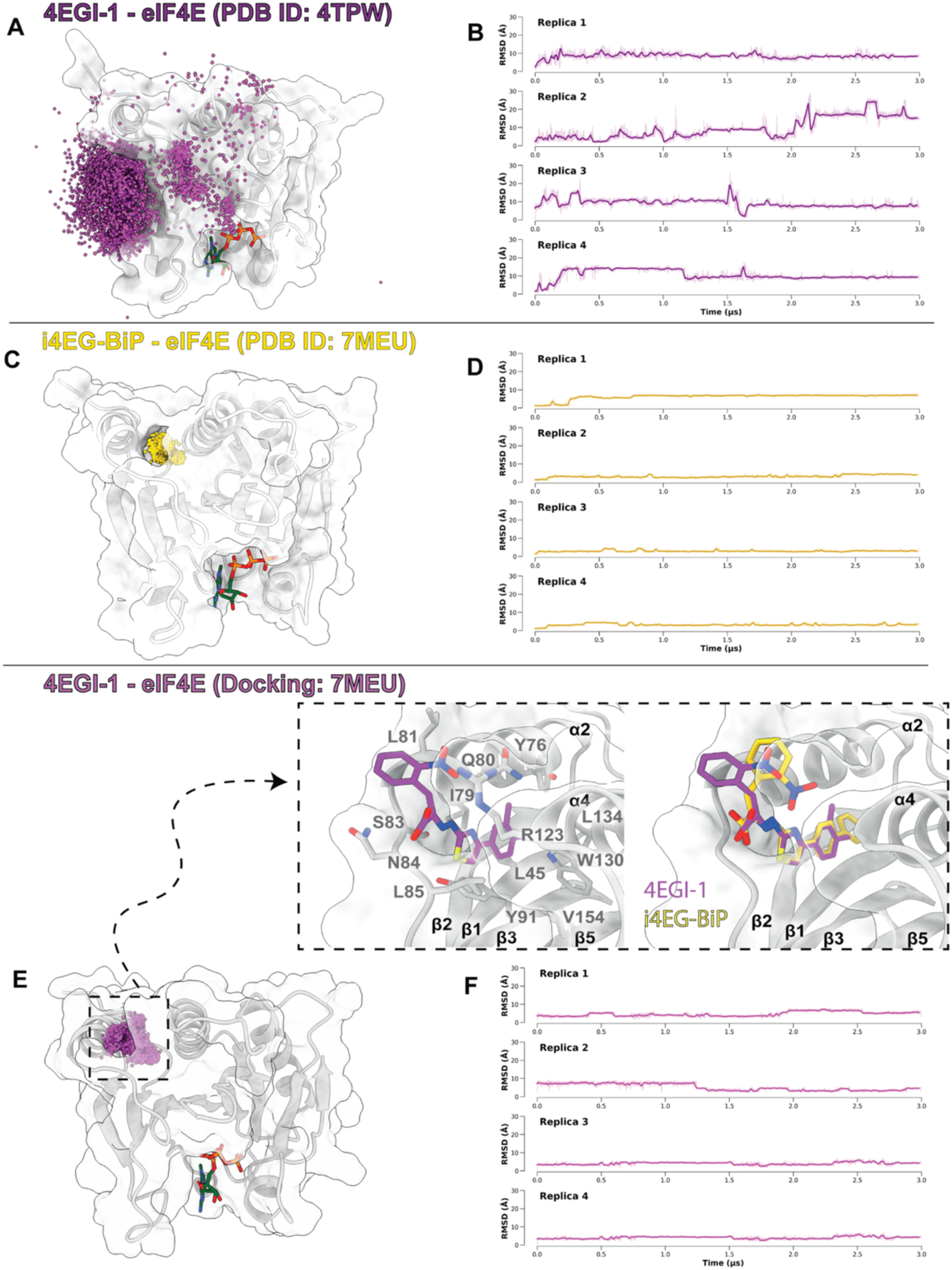
Time evolution of the crystallographic and docking poses of 4EGI-1 and i4EG-BiP bound to eIF4E. Ligand centers of mass (CoMs), sampled every 1 ns along the MD trajectories, are shown as magenta spheres for 4EGI-1 (A: crystallographic pose, PDB ID: 4TPW; E: docking pose) and yellow spheres for i4EG-BiP (C: crystallographic pose, PDB ID: 7MEU), respectively. Atomic details of the docked conformation of 4EGI-1 (magenta sticks) at the frontal binding site, overlaid with the crystallographic pose (PDB ID: 7MEU) of i4EG-BiP (yellow sticks), are shown in the inset of panel E. eIF4E is represented as silver ribbons with a transparent surface; m^7^GTP is displayed as green sticks. Key interacting residues are shown as sticks. Non-polar hydrogens are omitted for clarity. (B, D, F): Time evolution of ligands’ heavy-atom RMSDs across four independent replicas, with a 5 ns rolling average (bold lines) and raw data (faded lines).

Given these findings and the significant chemical similarity between the two compounds, we postulated that 4EGI-1 might adopt an alternative binding pose, resembling the frontal one observed with i4EG-BiP. To verify this hypothesis, we initially performed molecular docking calculations of 4EGI-1 at the i4EG-BiP’s binding cleft in eIF4E (PDB ID: 7MEU). Notably, docking results converged toward a single cluster of poses in which 4EGI-1 assumes an interaction pattern similar to that of i4EG-BiP, as shown by the superposition of its lowest energy docking pose and the crystallographic conformation of the latter (Figure 2E). To validate these results, we assessed the energetics and dynamics of the predicted docking mode through four independent 3 μs-long MD simulations. Interestingly, when starting from this alternative conformation, 4EGI-1 displays higher stability than in simulations initialized from the crystallographic lateral pose. Indeed, modest ligand’s CoM movement (Figure 2E) and RMSD fluctuations (Figure 2F) are observed across all replicas, suggesting that no significant rearrangement takes place. This emerging similarity in interaction fingerprints supports a shared preference for the frontal binding mode, however warranting further theoretical and experimental validation.

### Mutational Analysis Reveals Comparable Ligand Binding Profiles for 4EGI-1 and i4EG-BiP

To investigate the functional contribution of the two experimentally reported binding pockets of eIF4E, we designed a focused alanine-scanning mutagenesis guided by three independent criteria: (i) ligand–protein contact maps derived from X-ray structures and MD simulations; (ii) evolutionary conservation of binding-site residues (via ConSurf analysis)^47,48^; and (iii) evidence from previous mutational studies.

In the lateral binding pocket, key contact residues were identified by inspecting the crystal structure of the eIF4E/4EGI-1 complex (PDB ID: 4TPW) (Figure 3A). MD simulation data were excluded from this analysis due to the observed instability of the ligand in this region. The examination highlighted K49, R61, I63, H78, I79, and S83 as key contact residues. To avoid potential cross-talk with the adjacent frontal pocket, H78, I79, and S83 were excluded. As a result, K49A, R61A and I63A were prioritized for testing. Notably, the K49A mutant has been previously characterized in the literature^40^, and was therefore also used as an internal control.

**Figure 3.**
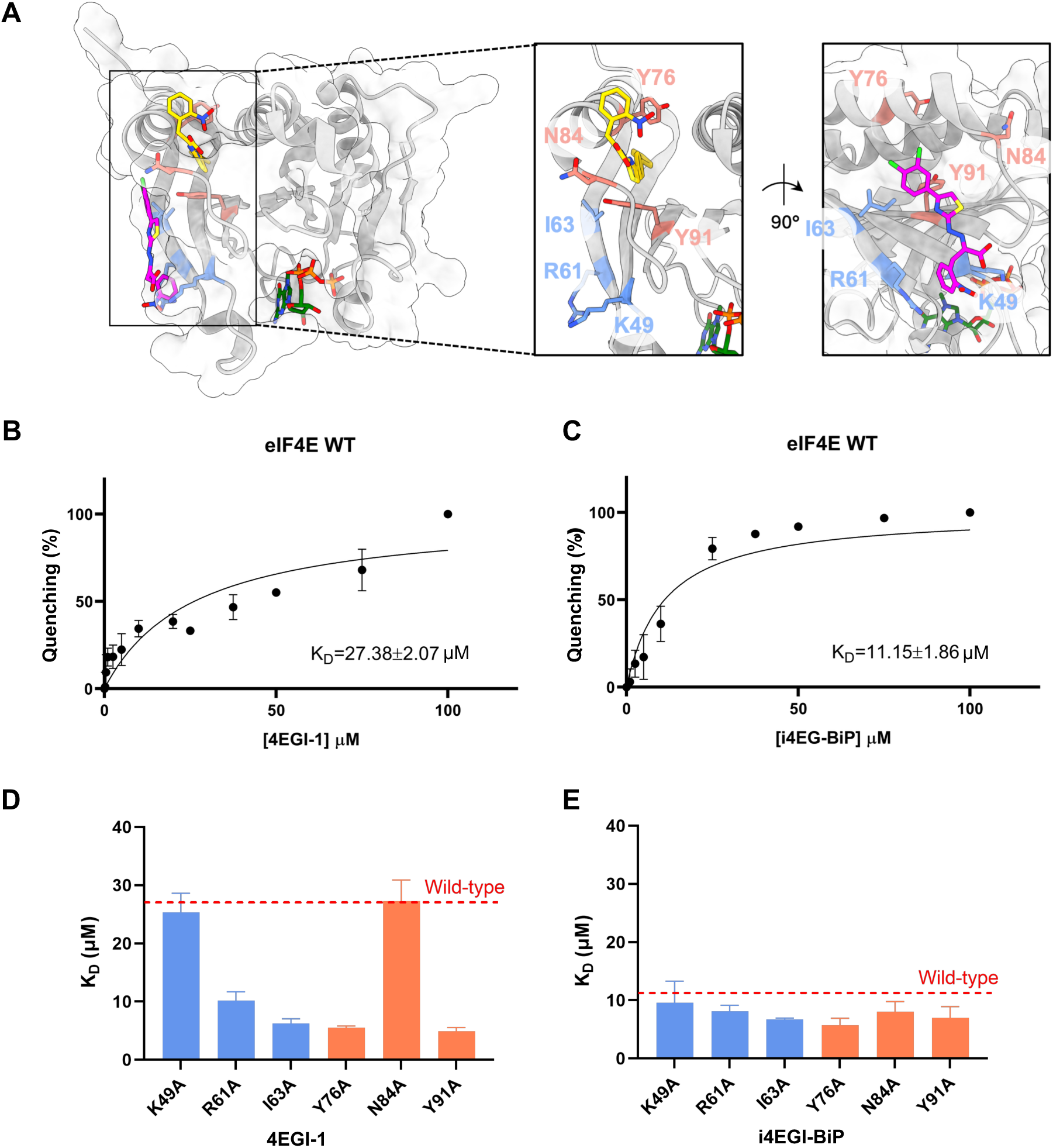
eIF4E Mutants Design and Testing. A) Mutated residues in lateral and frontal binding pocket of eIF4E, shown as insets. The protein is shown as light grey transparent surface and cartoon. 4EGI-1, i4EG-BiP and m^7^GTP are depicted as magenta, yellow and green sticks, respectively. Selected amino acids in the lateral and frontal binding pocket are depicted as blue and salmon sticks, respectively. B-C) Binding affinity measurements of 4EGI-1 (B) and i4EG-BiP (C) toward WT-eIF4E as determined by three independent tryptophan fluorescence quenching assays. Data are shown as percentage of intrinsic fluorescence intensity quenching versus inhibitor concentration and were fitted using a one-site specific binding model. Mean *K*_D_ ± standard deviation (SD) values are reported in the graph. D-E) Comparison of binding affinities across eIF4E mutants. Bar plots report the mean *K*_D_ value measured for each mutant. The WT-eIF4E *K*_D_ is shown as a red dotted reference line. Panel (D) shows *K*_D_ values for 4EGI-1 binding, and panel (E) shows *K*_D_ values for i4EG-BiP binding. Data represent the mean of three independent experiments; error bars represent SD.

In the frontal binding pocket, we examined the key ligand–protein interactions observed in the X-ray structure of i4EG-BiP (PDB ID: 7MEU) and in the docked conformation of 4EGI-1 within the same region. Structural inspection was then integrated with MD-based contact analysis, which revealed a modular binding architecture where the core and tail regions of both ligands remain stably anchored within the lipophilic tunnel between helices α2 and α4, while the solvent-exposed head adopts multiple orientations (Figure S1). This modular behavior provided a rationale to map interactions across the head, core, and tail moieties of each ligand (Figure S2) and prioritize mutation sites accordingly. Our analysis identified the following residues as main interaction partners: N84, L85, and R123 primarily contacting the head; Y91 the core; and Y76, I79, L93, W130, and L134 the tail. All leucine residues (L45, L85, L93 and L134) were discarded based on the typically mild effect of leucine-to-alanine substitutions^49,50^, and in the case of L85 and L93, also due to low contact persistence. Among the remaining candidates, N84 was chosen for the head region based on its higher contact frequency compared to R123. Y91, the only stably interacting residue within the core, was selected as sole representative for this region. For the tail, I79 was excluded to minimize cross-pocket effects, and W130 was discarded due to its high evolutionary conservation (Table S1), which raised concerns of global destabilization^51,52^. Thus, N84, Y91, and Y76 were retained as selective probes for the head, core, and tail regions, respectively^40,41^. Based on these criteria, we selected a panel of six single-point mutants targeting either the lateral (K49A, R61A, I63A) or the frontal (Y76A, N84A, Y91A) binding site, along with two double mutants (K49A–R61A and Y76A–L134A) to explore potential cooperative effects. Although the L134A substitution alone was predicted to exert minimal impact, it was considered for double mutations to evaluate possible synergy with nearby substitutions. All variants were first validated in silico to ensure structural integrity, showing no signs of instability in the apo state (Figure S3).

This curated set was then tested experimentally to assess the impact of individual binding-site perturbations on ligand recognition, with the goal of distinguishing site-specific contributions to 4EGI-1 and i4EG-BiP affinity. Wild-type (WT) and mutant forms of eIF4E were expressed in *E. coli* and purified (Figure S4A). The cap-binding activity of all variants was confirmed via m⁷GTP affinity pull-down assays (Figure S4B), and their binding affinities for 4EGI-1 and i4EG-BiP were determined using intrinsic fluorescence spectroscopy. First, we computed the dissociation constants (*K*_D_) of both inhibitors for wild-type eIF4E (Figure 3B-C), yielding values of 27.38 ± 2.07 μM for 4EGI-1 and 11.15 ± 1.86 μM for i4EG-BiP, consistent with previously reported IC_50_s (56.8 ± 1 μM for 4EGI-1 and 67.5 ± 2 μM for i4EG-BiP)^40,44^. This represents the first direct comparison of their *K*_D_ values under identical experimental conditions. Notably, i4EG-BiP exhibits a slightly lower *K*_D_ (Figure 3C), indicating higher affinity for eIF4E, in line with its superior performance in cellular assays^41^. Although the absolute changes in *K*_D_ are moderate, most point mutations, distributed across both the lateral and frontal pockets, elicited detectable shifts in affinity for at least one of the two ligands. This widespread sensitivity suggests that both regions contribute to ligand accommodation, and that neither compound relies on a single dominant hotspot but rather engages a distributed interaction network. Among the tested mutations, the largest differences between 4EGI-1 and i4EG-BiP were observed for K49A and N84A. Notably, both wild-type residues interact with the polar head of the ligands in the lateral and the frontal binding pose, respectively. In our simulations, this moiety exhibits the highest conformational variability and forms the least conserved contacts across the MD trajectories (Figure S1). These transient or partially populated interactions may underlie the more variable *K*_D_ shifts observed experimentally. In contrast, the hydrophobic core of the ligands acts as a stable anchoring motif across all trajectories (Figure S1). Consistent with this interpretation, the most pronounced variations in affinity, particularly for 4EGI-1, occur upon mutation of I63A, Y76A, and Y91A (Figure 3D and Figure S5A), three bulky and hydrophobic or aromatic residues involved in the accommodation of the ligand’s hydrophobic core within the two binding clefts. To further dissect site-specific contributions, we next examined the double mutants K49A–R61A (lateral pocket), and Y76A–L134A (frontal pocket). Neither double mutant significantly altered ligand binding (Figure S6), suggesting additive or compensable effects within each region.

Altogether, the overall impact of the mutations on binding affinity is modest yet notably consistent across the two ligands. In contrast to earlier studies reporting variations of up to one order of magnitude^40^, our measurements reveal more contained shifts, with similar trends observed for both compounds. Remarkably, the residues most affecting 4EGI-1 binding include two from the lateral pocket (I63 and R61) and two in the frontal site (Y76 and Y91). This cross-site sensitivity support, in line with our MD simulations, overlapping interaction profiles for 4EGI-1 and i4EG-BiP and point to coexisting binding modes, in which specific residues from both binding interfaces contribute to ligand affinity.

### Free Energy Simulations Corroborate the Frontal Ligand Binding Mode

MD simulations and mutagenesis data provided important insights into the binding behavior of 4EGI-1 and i4EG-BiP but were not sufficient to fully resolve their recognition mechanism. To achieve a deeper atomistic understanding, we performed free energy calculations using funnel metadynamics (FM)^45,46^. This enhanced sampling methodology has been extensively validated across diverse receptor classes^53–58^, and is now considered a state-of-the-art method for mechanistic studies of ligand binding. In FM, a history-dependent bias potential is added through well-tempered metadynamics (WT-MetaD)^59,60^ along carefully chosen collective variables (CVs), while a funnel-shaped restraining potential confines the ligand near the binding site. This setup enables multiple recrossing events between bound and unbound states and accurate reconstruction of the binding free energy surface (BFES).

In our implementation, we employed two CVs defined as the RMSD of the ligand with respect to the frontal and the lateral pose. This choice of CVs not only ensured proper sampling and energetic evaluation of these specific binding modes but also facilitated the exploration of possible alternative poses and the unbound state within the MetaD framework. Using this setting, both the simulations fully converged between 1.3 and 1.8 μs, as indicated by the stabilization of the BFES profile and the diffusive behavior of the employed CVs within this timescale interval (Figure S7 and S8). Importantly, this type of calculation also enabled us to estimate the absolute binding free energy of the ligands, computed as the difference between the bound and unbound states (see Methods for details). The resulting standard absolute binding free energy (Δ*G*^0^_b_) values are –9.93 ± 2.13 and – 11.4 ± 0.7 kcal mol^-1^ for 4EGI-1 and i4EG-BiP, respectively. Although this estimate carries the typical uncertainty associated with classical force fields, it is consistent with both the experimentally low-micromolar *K*_D_ of the ligands and the higher eIF4E affinity of i4EG-BiP, supporting the robustness and reliability of our computational approach.

The BFES of 4EGI-1, shown in Figure 4, reveals a clear global minimum at the frontal binding pose (state A), indicating it as the most thermodynamically favorable state. In contrast, the lateral pose (state B) appears to be a higher-energy, less stable alternative state. These results are consistent with unbiased MD simulations, in which ligand instability is systematically observed when initialized from the lateral pose. Conversely, the frontal pose demonstrates robust stability across all simulation replicas, reinforcing its role as the preferred binding conformation not only for the i4EG-BiP inhibitor but also for 4EGI-1. Notably, no additional low-energy basin is detectable in the BFES, thereby excluding any further alternative ligand-receptor interaction mode.

**Figure 4.**
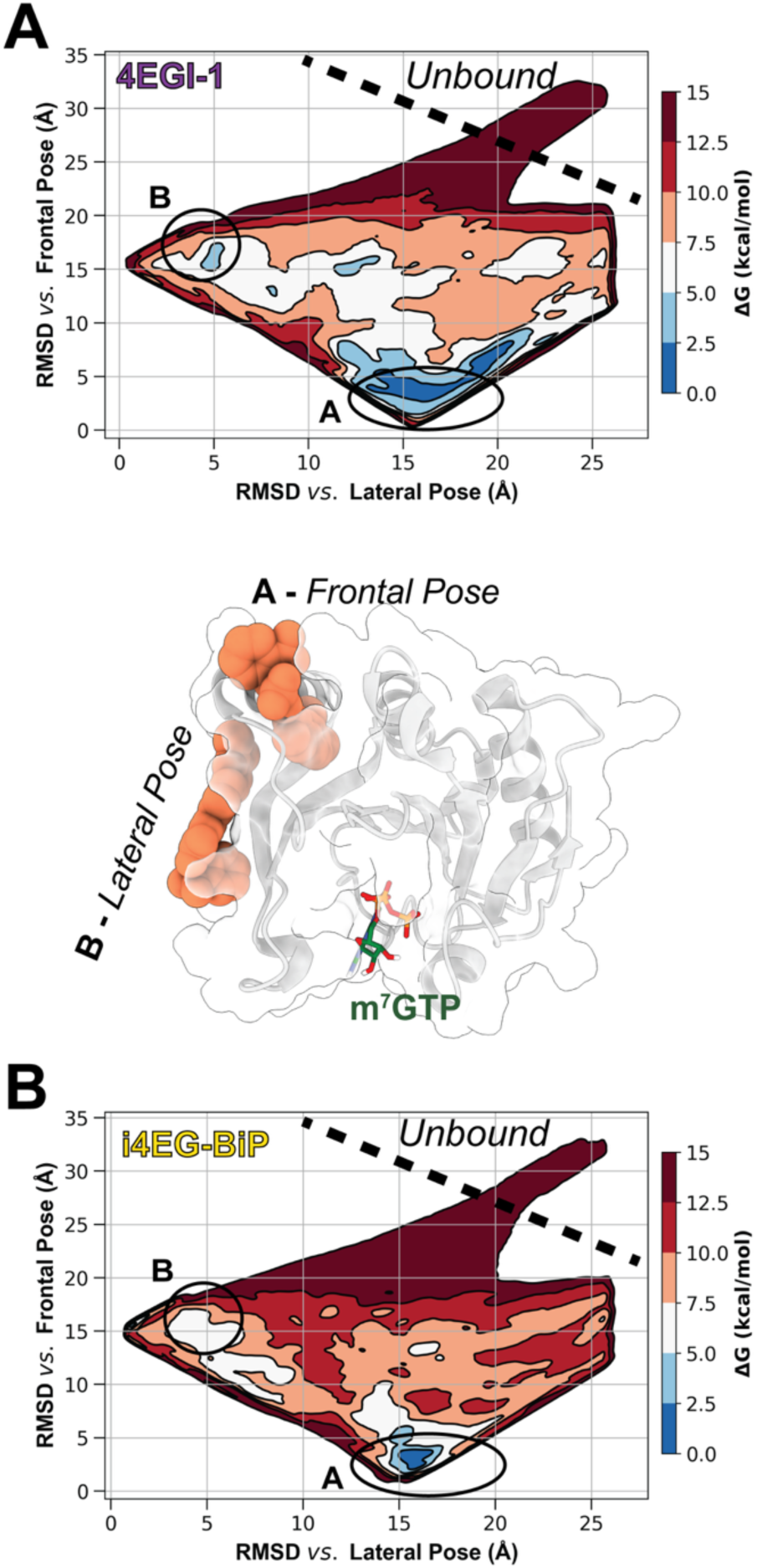
Binding Free Energy Calculations with Funnel Metadynamics. Binding free energy surface (BFES) of 4EGI-1 (A) and i4EG-BiP (B) at eIF4E, projected along the RMSD relative to the lateral (x-axis), and to the frontal (y-axis) poses. Contour lines (isolines) are plotted at 2.5 kcal/mol intervals. Energy basins A and B correspond to the frontal and lateral binding modes, respectively. Representative structures of each ligand pose are shown in the middle panel as salmon spheres; the protein is depicted as silver cartoon and transparent surface.

A similar energetic profile is obtained for i4EG-BiP (Figure 4), which displays an even more pronounced preference (approximately 2 kcal/mol) for the frontal pose over the lateral one. This stronger stabilization is consistent with the enhanced robustness observed in unbiased MD trajectories, where the frontal binding mode of i4EG-BiP remains even more persistently engaged throughout the simulations. Altogether, these results support a converging picture from enhanced sampling and unbiased simulations, corroborating the frontal pose as the dominant and functionally relevant binding mode for both ligands. Conversely, the lateral pose emerges as a higher-energy state in all cases, yet is relatively more accessible for 4EGI-1, suggesting that it may act as an alternative binding mode or represent a transient intermediate sampled along the recognition pathway.

### Ligand-Induced Allosteric Antagonism Shapes Partner-Motif Recognition at eIF4E

Once established the binding behaviors of 4EGI-1 and i4EG-BiP, we investigated how these ligands modulate eIF4E’s interactions with its physiological partners, disrupting eIF4G binding while promoting 4E-BP association. Structural data show that these partner proteins interact with eIF4E through three distinct motifs: (i) a conserved canonical motif (CM, YXXXXLΦ) that binds the dorsal surface in a helical arrangement; (ii) a more variable, flexible non-canonical motif (NCM) targeting the lateral surface; and (iii) a linker region connecting the two motifs in an elbow-shaped configuration (Figure 5A and Figure S9)^9,26^. When the lateral binding mode of 4EGI-1 was first reported, structural information on NCM–eIF4E recognition was still unavailable. This led to the initial hypothesis that ligand binding at the lateral site of eIF4E might trigger long-range allosteric changes at the dorsal site, disrupting the recruitment of partner proteins via their CM^39,40^. Subsequent structural studies, however, revealed a substantial overlap between the NCM binding site and the lateral ligand pocket, supporting a model of direct competition between the ligand and any eIF4E partner protein^9^. Yet this interpretation conflicted with experimental findings showing that 4EGI-1 selectively impairs eIF4G recruitment while enhancing 4E-BP binding ^39,43^. Furthermore, comparable biochemical effects have been reported for i4EG-BiP, and more recently, other ligands that engage the frontal pocket ^32,41^, suggesting that a shared allosteric mechanism underlies their functional outcome. In line with this evidence, both 4EGI-1 and i4EG-BiP induce a characteristic conformational rearrangement in eIF4E observed neither in the apo nor partner-bound states. This transition involves the unfolding of a short 3_10_-helix (residues S82–L85) and a one-turn extension of the adjacent α2-helix (residues H78–S82), which together expose the frontal pocket otherwise occluded in the apo form. We refer to this conformation as the “extended” state^40,41^, in contrast to the “reduced” conformation of the apo structure^9,39^ (Figure 5B). Conversely, engagement of eIF4G- or 4E-BP1-derived peptides stabilizes the “reduced” state of eIF4E, preserving the 3_10_-helix and enabling NCM binding at the lateral surface. This conformation is sterically incompatible with both lateral ligand binding and the structural transition required to access the frontal pocket, suggesting an allosteric antagonism between partner association and ligand-induced rearrangement.

**Figure 5.**
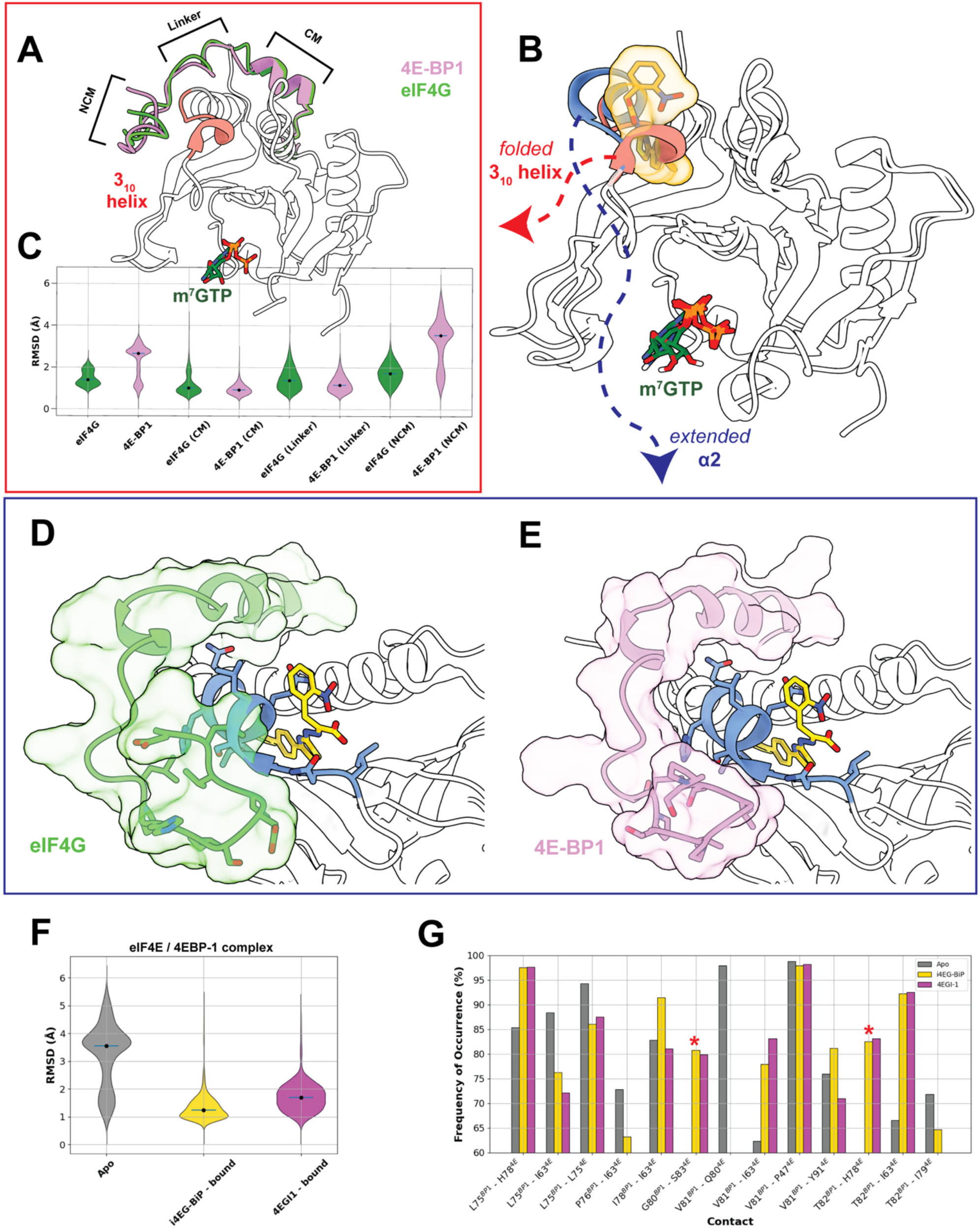
Analysis of eIF4G and 4E-BP1 peptides bound to eIF4E, with and without 4EGI-1 and i4EG-BiP. A) Tridimensional representation of the interaction between eIF4E (white transparent cartoon) and its peptide partners: eIF4G (green cartoon, PDB ID: 5T46) and 4E-BP1 (pink cartoon, PDB ID: 5BXV). B) Conformational rearrangement of eIF4E (white transparent cartoon) upon ligand binding, involving the unfolding of the 3_10_-helix (residues S82–L85) and the one-turn extension of the adjacent α2-helix (residues H78–S82). The ligand shown is i4EG-BiP (yellow transparent surface and stick), provided as a representative example. m^7^GTP is depicted as green sticks. C) Violin plots illustrating the RMSD distributions (Cα atoms) for the complete peptides and their eIF4E-binding motifs throughout the MD simulations. D-E) Superposition of the most representative MD-derived conformations of the eIF4G (green cartoon and transparent surface) and 4E-BP1 (pink cartoon and transparent surface) peptides, in complex with eIF4E (white transparent cartoon), onto the target protein structure (PDB ID: 7MEU) bound to i4EG-BiP (yellow sticks). The two conformations were obtained through a cluster analysis with a cut-off of 1.5 Å. F) Violin plots showing the RMSD of the 4E-BP1 NCM (Cα atoms) over the MD trajectories of the eIF4E/4E-BP1 complex, either in the absence or presence of i4EG-BiP or 4EGI-1 (frontal pose). G) Frequence of occurrence for the most persistent (≥ 60%) contacts between 4E-BP1 (NCM) and eIF4E contacts observed during MD simulations, either in the absence or presence of i4EG-BiP or 4EGI-1 (frontal pose).

To clarify this mechanism, we investigated the dynamic interplay between eIF4E, its protein partners, and the two ligands, focusing on the frontal binding mode that here emerged as the most stable configuration. First, we performed extensive MD simulations of eIF4E in complex with co-crystallized peptides from eIF4G (PDB ID: 5T46^9^) and 4E-BP1 (PDB ID: 5BXV^39^), each in four independent 3-μs replicas. Global RMSD analysis of the peptide backbones throughout these simulations show greater stability of the eIF4G-derived fragment compared to 4E-BP1 (Figure 5C). To refine this observation, we calculated individual RMSDs for the CM (residues 50–63), linker (residues 64–72), and NCM (residues 73–84) regions (Figure 5C). Both the CM and linker of the two peptides remained stably associated with eIF4E across all simulations, as also reflected by the persistence of native contacts involving these domains (Tables S2 and S3). Conversely, the NCM of the 4E-BP1 peptide shows higher flexibility compared to that of the eIF4G-derived fragment (Figure 5C and Figure S10), consistent with previous reports on 4E-BP2^61^. This increased mobility is mirrored by amplified fluctuations in the interacting region of eIF4E (residues 77-85, Figure S11). These differences likely arise from two stabilizing contacts formed by L641 of the eIF4G peptide with H78 and Q80 of eIF4E (Figure S10), which are absent in the 4E-BP1 complex. Cluster analysis across the MD trajectories further supports this interpretation, showing that the NCM of the eIF4G peptide consistently adopts a single dominant conformation, indicative of a rigid and well-defined structural preference. In contrast, the NCM of 4E-BP1 populated multiple clusters, reflecting its greater conformational flexibility.

### Frontal-Ligand Binding Enables Ternary-Complex Formation and Stabilization

Building on this framework, we next assessed whether the sampled conformations of eIF4G- and 4E-BP–derived peptides were compatible with the ligand-bound extended state of eIF4E, thereby enabling formation of a ternary complex. To this aim, we superimposed the dominant conformers from cluster analysis onto the crystal structure of eIF4E bound to i4EG-BiP (PDB ID: 7MEU), focusing on the NCM region. This comparison showed that the rigid NCM conformation of the eIF4G-derived peptide is incompatible with the structural rearrangement induced by ligand binding, namely the unfolding of the 3_10_-helix and the extension of α2 (Figure 5D). This likely precludes access to the frontal pocket, explaining the mutual exclusivity of ligand and eIF4G binding. Conversely, the more flexible NCM of 4E-BP1 can better adapt to these structural changes (Figure 5E), allowing for co-binding and even enhanced association, in agreement with experimental data. On the other hand, both peptides showed steric clashes with ligands bound at the lateral pocket, suggesting that only frontal binding supports ternary complex formation. These results thus support a model in which frontal, but not lateral, ligand binding permits or enhances peptide association depending on their flexibility. To validate this hypothesis, we conducted additional MD simulations (4 replicas of 3 μs each) of eIF4E in its extended conformation complexed with the 4E-BP1 peptide and either 4EGI-1 or i4EG-BiP.

In both the ternary systems, the 4E-BP1 NCM exhibits reduced RMSD fluctuations relative to the binary complex, indicating enhanced stability in the presence of ligands (Figure 5F). This effect can be primarily attributed to new interactions, specifically G80^BP1^-S83^4E^, T82^BP1^-78^4E^, formed between the 4E-BP1 peptide and the additional helical turn (residues N77 to S85) unique to the “extended” eIF4E conformation. Several pre-existing contacts are also reinforced, including L75^BP1^-H78^4E^, V81^BP1^-I63^4E^, T82 ^BP1^-I63^4E^ (Figure 5G and Figure S12A). These contributions outweigh the minor loss of the V81^BP1^-Q80^4E^ interaction and a slight reduction in the frequency of the L75^BP1^-I63^4E^ and P76^BP1^-I63^4E^ contacts (Figure S12B). Concurrently, the binding poses of both ligands also display increased stability in the presence of 4E-BP1 relative to their already steady behavior in binary complexes (Figure S13). Taken together, these results strongly support a cooperative mechanism in which ligand engagement at the frontal pocket promotes the extended eIF4E conformation, thereby favoring 4E-BPs association and disfavoring eIF4G binding, an outcome that is incompatible with the “reduced” state.

## DISCUSSION

Protein translation represents a critical process in cancer cells, which are highly dependent on the rapid production of oncogenic proteins to survive and adapt to environmental stress ^12–16^. In this context, targeting translation initiation has long been recognized as a promising anticancer strategy ^1,18,19,22–27^. Among initiation factors, eIF4E stands out as a particularly attractive target, as it serves as the rate-limiting component of cap-dependent protein synthesis and acts as a convergence point for multiple key oncogenic signaling pathways^7,8^. Despite substantial progress in developing small-molecule inhibitors that interfere with upstream kinases, disrupt protein-protein interactions^29–31^, or prevent binding to capped mRNA^35,36^, the translation of these tool compounds into clinically effective inhibitors has proven challenging. This is mainly because eIF4E is an inherently difficult target, being small and hydrophobic, and prone to forming protein-protein interactions. Among the various strategies for blocking eIF4E, inhibiting its binding to eIF4G appears to be the most direct and promising^1,30–32,39–41^. To date, the best characterized small molecules that interfere with eIF4E/eIF4G interaction are 4EGI-1^40,43^ and its analogue i4EG-BiP^41^, which were proven effective to inhibit protein synthesis and cell proliferation. Using intrinsic fluorescence quenching assays, we provide here the first direct comparison of their binding affinities for wild-type eIF4E under identical experimental conditions, yielding *K*_D_ values of 27.38 ± 2.07 μM for 4EGI-1 and 11.15 ± 1.86 μM for i4EG-BiP. Despite sharing the same functional outcomes, static X-ray data suggest that the two ligands engage eIF4E through two distinct binding modes, with 4EGI-1 interacting at the lateral pocket^40^ and i4EG-BiP binding to the frontal site^41^. This divergence is unusual for structurally related ligands, yet converging on similar regulatory effects. Through extensive molecular simulations and free energy calculations, our work reveals the frontal binding mode is thermodynamically favoured for both 4EGI-1 and i4EG-BiP, whereas the lateral pose of 4EGI-1 likely represents a metastable intermediate along its recognition pathway. Computational predictions are supported by site-directed mutagenesis, in which six residues identified as critical for the frontal (K49, R61, I63) or lateral (Y76, N84, Y91) binding modes were mutated to alanine, individually or in pairs (K49A–R61A and Y76A–L134A). Binding assays on mutants show similar trends for both ligands, with most substitutions having minor effects, whereas only I63A, Y76A, and Y91A showing modest *K*_D_ reduction, particularly in the case of 4EGI-1. These results would suggest that alanine substitutions at certain positions do not necessarily disrupt binding through loss of critical contacts, but could modulate the interaction through indirect effects such as reducing steric hindrance or altering local electrostatics. The lack of a single mutation with consistent disruptive effects, combined with the similar impact observed across various substitutions, supports the view that 4EGI-1 and i4EG-BiP share the binding mechanism, and that ligand recognition by eIF4E is inherently dynamic. It is worth noting that other studies have investigated eIF4E mutations in both the lateral and frontal pockets, but primarily in relation to their impact on eIF4E activity, such as eIF4G displacement^40^ or proliferation rescue assays in cancer cells^32^, and not in terms of direct ligand binding evaluation. Our findings provide complementary molecular insight by dissecting the thermodynamic contributions of such residues to ligand recognition, thereby bridging the gap between structural modeling and functional outcome. A central advance of this study is the clarification of how 4EGI-1 and i4EG-BiP modulate the selectivity of eIF4E for its physiological partners, disrupting interaction with eIF4G while reinforcing 4E-BP association. Previous structural and biochemical data offer seemingly contradictory insights into the underlying mechanism^1,9,32,39,40^. On one hand, X-ray studies suggest that ligand binding at the lateral pocket could sterically hinder partner recognition through their non-canonical motifs (NCMs)^9^. On the other, they show that ligand binding at the frontal pocket induces a rearrangement of eIF4E involving unfolding of the 3_10_-helix and extension of the α2-helix^40,41^. This structural transition is apparently incompatible with the binding of either eIF4G or 4E-BPs, raising questions about how ligand binding could enhance 4E-BP recruitment.

Our work resolve this contradiction by showing that the ligand-induced rearrangement, while sterically incompatible with eIF4G, is in fact highly permissive, and even favourable, for 4E-BP1 binding. This difference stems from the distinct structural properties of their respective NCMs. In particular, the rigid NCM of eIF4G adopts a well-defined conformation that clashes with the extended state of eIF4E, effectively precluding frontal pocket access. In contrast, 4E-BP1 exhibits a more flexible NCM capable of adapting to the rearranged α2/loop segment, thereby forming new stabilizing contacts in the ternary complex.

Molecular dynamics simulations of the ternary complexes of eIF4E with a 4E-BP1 peptide and either 4EGI-1 or i4EG-BiP, confirm this cooperative effect. In the presence of either ligand, the 4E-BP1 peptide exhibits reduced structural fluctuations and an expanded interaction network with eIF4E. Simultaneously, the ligands themselves adopt more stable binding poses. These findings provide a mechanistic explanation for the observed mutual exclusivity of ligand and eIF4G binding, and the concomitant enhancement of 4E-BP1 association. Crucially, no such stabilization is possible from the lateral pose, which is sterically incompatible with ternary complex formation. Therefore, productive ternary assembly strictly requires the frontal pose, identifying this configuration as the functionally active binding mode of both inhibitors.

Overall, this study illustrates how ligand-induced conformational plasticity can be harnessed to reprogram the interaction landscape of a highly networked regulatory hub such as eIF4E. By revealing how specific ligands reshape protein-protein interaction selectivity, our findings offer a conceptual framework for structure-guided design of small molecules that not only block individual interactions but exploit allosteric cooperativity to selectively stabilize desired complexes. This may enable a new class of translational modulators with applications in cancer and other diseases.

## MATERIALS AND METHODS

### Molecular Docking

For docking calculations of 4EGI-1 at the frontal binding site, we selected the X-ray structure of eIF4E bound to i4EG-BiP (PDB ID: 7MEU^41^), which was prepared using the Protein Preparation Wizard in Maestro v. 2020-4^62^. Missing hydrogen atoms were added. Protonation states and hydrogen bonding networks at local pH were assigned using the PropKa program^63^ included in Maestro^64^. Finally, the positions of all the hydrogens were minimized, using the OPLS-2005 force field^62^. The tridimensional structure of 4EGI-1 was retrieved from the X-ray structure of the ligand in complex with eIF4E (PDB ID: 4TPW^40^) and then prepared using LigPrep. The ligand’s 3D tautomeric and protonation state in the pH range of 7.0 ± 1.5 were, however, assigned using Epik^65^. Docking calculations were performed with the grid-based program Glide^66–68^. The Receptor Grid Generation tool was used to generate a grid box of 20 Å × 20 Å × 15 Å centered on the i4EG-BiP’s binding pocket. Docking was performed in the standard precision (SP) mode^67,68^ using the OPLS-2005 force field^62^. Otherwise, default parameters were applied. Docking results showed clear convergence toward a single binding mode, with the best-scoring pose yielding a docking score of –6.6.

### Ligand Parametrization

For the organic compounds 4EGI-1 and i4EG-BiP, atom types, and all bonded and non-bonded parameters were assigned using the Generalized AMBER Force Field (GAFF2)^69^. For m^7^GTP, atom types and parameters were primarily taken from the Amber RNA force field (OL3), whereas missing parameters were derived from GAFF2 to ensure compatibility with the Amber14SB^70^ force field used for the protein. Geometry optimizations were performed using Gaussian16 (Rev. A03)^69^ in a two-step procedure at the Hartree-Fock (HF) level of theory. The first step employed the 3-21G basis set, followed by a second optimization using the 6-31G* basis set. During the latter step, the electrostatic potential (ESP) was also computed and used to derive atomic partial charges via the two-stage Restrained Electrostatic Potential (RESP) fitting procedure implemented in Antechamber^71^. Both 4EGI-1 and i4EG-BiP contain two dihedral angles involving conjugated systems, which often require careful parametrization within the AMBER force field^72,73^. Thus, we conducted a detailed quantum mechanical (QM) analysis of these torsions, located between: (i) the thiazole and benzene rings, and (ii) the thiazole ring and the adjacent hydrazine group. Initial QM torsional scans were performed in vacuo using Gaussian16. Each dihedral angle was fixed at sequential values while the rest of the molecule was optimized (Figure S14). Calculations were carried out at the MP2/6-31G* level of theory, following the standard GAFF protocol.

The resulting QM potential energy profiles were then compared to those predicted by GAFF2. While the GAFF2 profile for the thiazole–benzene dihedral showed good agreement with the QM reference, significant discrepancies were observed for the torsion between the thiazole ring and the adjacent hydrazine group. To correct this, we derived custom torsional parameters that better reproduced the QM potential energy surface, using the standard AMBER dihedral functional form: *E_dih_* = *k* (1 + cos (*ný* − *ψ*)), where k is the force constant, n is the periodicity, *ϕ* is the torsion angle, and *ψ* is the phase angle. The optimal parameter set was obtained using an in-house genetic algorithm and involved a combination of two torsional terms. The final fitted expression was: *E_dih_* = −1.25 (1 + cos (*ý* − *π*)) + 2.2 (1 + cos (*ý - π*)). This function accurately captured the behavior of the QM energy profile and was consequently used in MD simulations.

### Molecular Dynamics Simulations

The simulated systems were retrieved from the PDB, generated through docking calculations or manually assembled, as detailed in Table S4. All proteins were first pre-processed using Protein Preparation Toolkit implemented in Maestro^64^. Protonation states were assigned using Epik^65^ at pH 7.0, and N- and C-terms were capped. The topology files for the systems were obtained using the tleap program from AmberTools2018^74^ and then converted to the GROMACS format with ParmEd. eIF4E and the peptides derived from eIF4G and 4E-BP1 were parametrized using the ff14SB Amber Force Field^70^, while 4EGI-1, i4EG-BiP and m^7^GTP were parameterized as described above. Each system was solvated in a TIP3P^75^ cubic water box with a 12.0 Å edge distance (16.0 Å in funnel metadynamics calculations), and neutralized with Cl^-^ and Na^+^ ions modeled with Joung and Cheatham parameters^76^.

The GROMACS 2020.6 code^77–80^ was used to perform the simulations. A cutoff of 12 Å was applied for short-range interactions. Long-range electrostatic interactions were calculated using the particle mesh Ewald (PME) method^81^ with a 1.2 Å grid spacing in periodic boundary conditions (PBC). The non-iterative LINCS algorithm^82^ was employed to constrain bonds, enabling a 2-fs integration time step with the leapfrog algorithm.

To resolve all steric clashes, each system underwent 10,000 steps of steepest descent energy minimization in two phases. First, the system’s heavy atoms were fixed to relax only the hydrogens and water molecules; then, all atomic positions were minimized. Subsequently, each complex was equilibrated and heated to 300 K. Thermalization was performed in the canonical (NVT) ensemble over three steps of 0.5 ns each), gradually increasing the temperature to 100 K, 200 K, and 300 K using the Berendsen thermostat^83^, During this process, positional restraints were gradually reduced from 47.80 to 23.90 kcal/mol/nm^2^ and to 11.95 kcal/mol/nm^2^. These three steps were followed by two equilibration stages in the isothermal-isobaric (NPT) ensemble at 1 bar, using the V-rescale thermostat^84^ and the Parrinello-Rahman barostat^85^. During the first NPT step (1 ns), restraints of 5.97 kcal/mol/nm^2^ were applied and then gradually released over the second step (5 ns). The production phase followed seamlessly from the final equilibration step, maintaining the same thermostat and barostat settings. All calculations were carried out in four independent replicas of 3 microseconds each, initialized with random velocities.

All simulations were conducted in the presence of m^7^GTP, which remains stably bound to eIF4E in all systems (Figure S15). RMSD calculations, center of mass tracking, and cluster analysis were all performed using GROMACS tools^77^. Cluster analysis was carried out with the gromos algorithm^86,87^. Residue contact analysis for identifying mutation targets was performed in VMD using an in-house script^88^, while interactions between eIF4E and eIF4G or 4E-BP1 peptides were evaluated using the ProLIF tool^89^.

### Funnel Metadynamics

Funnel Metadynamics (FM) simulations were performed using GROMACS 2020.6^77–80^ patched with PLUMED 2.7.0^90–92^. A funnel-shaped restraining potential was applied to encompass both the lateral and frontal binding regions of the protein (Figure S16). The funnel potential was defined by the following parameters: (i) the height of the conical portion (*Zcc =* 28 Å), (ii) the aperture angle (α = 0.55 rad), and (iii) the radius of the cylindrical section (R_cyl_ = 1 Å). The binding free energy surface F(*s*,t) as a function of the selected collective variables (CVs) was computed using well-tempered metadynamics^59,60^, according to the following expression:

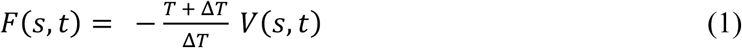

where *V(s,t)* is the bias potential added to the system, *T* is the temperature of the simulation, and Δ*T* is the fictitious temperature increase associated with the CVs. A bias factor of 20 was used, corresponding to Δ*T* = 5700 K. Gaussian hills with an initial height of 0.956 kcal/mol were deposited every 1 ps and gradually decreased over time, as per the well-tempered scheme.

As CVs, we employed the RMSD of the ligand heavy atoms with respect to: (i) the lateral and (ii) frontal binding poses (PDB IDs: 4TPW^40^ and 7MEU^41^, respectively). Prior to RMSD evaluation, the protein was aligned on the Cα atoms of its secondary structure elements to ensure consistent structural comparison.

In addition to the funnel restraint, the exploration was limited by upper walls set at 2.56 nm for the RMSD vs. lateral pose and 3.4 nm for the RMSD vs. frontal pose, to hamper nonphysical sampling beyond the relevant binding region.

The standard absolute binding free energy (Δ*G*^0^_b_) of the two compounds was computed according to the equation:

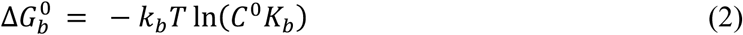

where *C^0^* = 1/1660 Å^−3^ is the standard concentration of 1 M for all reacting molecules, *k_B_* is the Boltzmann constant, and T is the temperature of the system (in Kelvin). The equilibrium binding constant *K_b_* is defined as:

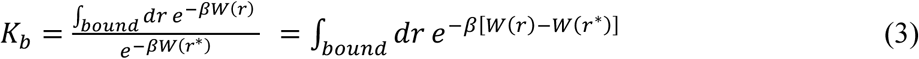

where *r* denotes positions along the chosen collective variable (CV) in the bound state, *W(r)* is the potential of mean force (PMF) at position *r* and *W(r*)* the PMF at a reference unbound state. The bound region was defined as the ensemble of configurations in which the value of the projection of the ligand’s CoM position onto the funnel axis (i.e., *fps.lp* CV^93^) was lower than 2.5 nm. The unbound state was chosen within the isoenergetic region of the cylindrical funnel section, where *fps.lp* exceeded 2.8 nm. Prior to Δ*G*^0^_b_ calculation, the free-energy profile along *fps.lp* was reconstructed using the bias reweighting procedure developed by Bonomi et al.^94^, as implemented in PLUMED. To assess convergence of the FM simulation, we first verified the quasi-diffusive behavior of the biased CVs (Figure S7A-E and S8A-E). Next, Δ*G*^0^_b_ was evaluated at increasing time intervals, showing convergence toward a plateau beginning at around 1.2 microseconds (Figure S7E). Error bars were computed as the standard deviation of the time-weighted mean within this convergence window, following the protocol described in ^45^. Finally, visual inspection of the binding free energy surface (BFES) confirmed that its global shape remained stable over the convergence period (Figure S7F and S8F).

### Production of Recombinant Wild-Type and Mutant eIF4E Proteins

Human full-length eIF4E cDNA was introduced into pMCSG7 plasmid and expressed in *Escherichia coli* BL21(DE3) pLysS. The recombinant protein, carrying an N-terminal 10×His tag, was recovered from inclusion bodies, denatured in 6 M guanidine hydrochloride, refolded, and purified in a buffer containing 50 mM HEPES pH 7.8, 300 mM KCl, 2 mM DTT, and 10% glycerol. For fluorescence quenching experiments, the protein was dialyzed against 20 mM HEPES pH 7.8 and 300 mM KCl, and stored at –80 °C. Detailed procedures for protein purification will be provided in a related manuscript currently in preparation. The protein was extracted from inclusion bodies using 6 M guanidine and then dialyzed against 20 mM HEPES pH 7.8, 300 mM KCl. Purified protein was stored at −80 °C until use. Single mutations (K49A, R61A, I63A, Y76A, N84A, Y91A) and double mutations (K49A-R61A, Y76A-L134A) were introduced through site-directed mutagenesis using primers carrying the desired nucleotide substitutions at their 5’ ends. The coding sequence of eIF4E was amplified with PCR and the parental DNA was digested using DpnI (NEB). The amplification product was treated with T4PNK followed by T4 ligase. All mutants were expressed and purified under the same conditions as for the wild type. Protein integrity and purity were confirmed by SDS-PAGE followed by Comassie Blue staining.

### Cap-binding Assay

Affinity pull-down assay of eIF4E wild-type and its mutants was performed using an Immobilized γ-aminophenyl-m^7^GTP (C_10_-spacer) resin (Jena Bioscience). Briefly, 30 µL of resin was pre-equilibrated in the protein storage buffer (20 mM Hepes pH 7.8, 300 mM KCl) and blocked with 0.1 mg/mL Bovin Serum Albumin (BSA) for 1 h at 4 °C under rotation to minimize any unspecific interactions. Following three washes with the same buffer, 10 µg of each protein was incubated with the resin for 2 h at 4°C under rotation. Bound proteins were eluted using 30 µL of 2X Laemmli Sample Buffer. Samples were analyzed by a 12.5% SDS-PAGE followed by Western Blot using an antibody against eIF4E (#9742 Cell Signaling Technology).

### Intrinsic Tryptophan Fluorescence Quenching Assay

The high intrinsic fluorescence of human eIF4E, attributed to its eight tryptophan residues, has been a useful feature in several prior quenching assays^39,40^. A mix solution containing 1 µM of proteins and increasing concentrations of either 4EGI-1 (324517, Sigma-Aldrich) or i4EG-BiP (HY-150903, ChemExpress) up to 100 μM or 150 μM of both inhibitors was prepared in analysis Buffer (20 mM Hepes pH 7.8, 300 mM KCl). Stock solutions of inhibitors were prepared in dimethylsulfoxide (DMSO) and serially diluted in the analysis buffer. Fluorescence spectra were acquired using a Tecan Infinite F Plex plate-reader, with 280 nm excitation and 340 nm emission filters. Fluorescence measurements were performed in 96-well plates (flat-bottom, black, Corning 3991) with a final volume of 150 µL per well. Raw fluorescence values were corrected by subtracting the background signal. For each dataset, the fluorescence in the absence of inhibitor (*V_max_*) and in the presence of the highest inhibitor concentration (*V_min_*) were identified and used as internal references. Fluorescence readings were then normalized by setting *V_min_* to 0 and *V_max_* to 100, respectively. Normalized fluorescence (*F_%_*) was calculated as:

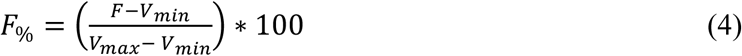

where *F* is the fluorescence value of each well. Percentage of intrinsic fluorescence intensity quenching was then calculated as:

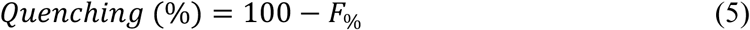

The resulting data were fitted in GraphPad Prism using non-linear regression with a one-site specific binding model, fixing *B_max_* fixed at 100. All experiments were performed in triplicate.

## ACKNOWLEDGMENTS

This article is in memory of Daniele Di Marino (1982-2024), who conceived the project and whose early insights significantly contributed to its development. We acknowledge funding support from the Italian Foundation for Cancer Research (AIRC, Project No. IG 807 2022 ID 27534) and the Italian Ministry of University and Research (MUR), PRIN 2022 (Project No. 202242MEP7). Computational resources were provided by the CINECA ISCRA B project “Dynamical and mechanistic insights into ligand binding to the translation initiation factor 4E (eIF4E)” (MD-eIF4E), which we gratefully acknowledge.

## AUTHOR CONTRIBUTIONS

D.D.M., A.Rm and F.S.D.L. and conceptualized the study. A.Rs. and V.M.D.A. performed computational studies. J.R. and G.R.B. expressed and purified mutant proteins and performed the fluorescence quenching experiments. A.Rm. optimized the protocols and supervised the experimental workflow. A.Rs., V.M.D.A. and F.S.D.L. reviewed, analyzed, and interpreted the computational data. A.L.T. and A.Rm. reviewed, analyzed, and interpreted the experimental data. All authors wrote, edited, and reviewed the manuscript. All living authors approved the final version of the manuscript.

## DATA AVAILABILITY

All the input files and trajectory data sets are published on a public Zenodo folder and freely available at the following link: 10.5281/zenodo.18233509.

## ADDITIONAL INFORMATION

- **Supplementary Material:** The online version contains supplementary material.

