## Supplementary file for "Ligand-Induced Structural Dynamics Drive Allosteric Regulation of Translation Initiation Factor eIF4E"

[†]Deceased. Contributed to the conception and early development of the project.

### Supplementary Information

| TABLE OF CONTENTS | |
|---|---|
| **Figure S1**: Conformational dynamics of the frontal binding mode of 4EGI-1 and i4EG-BiP during MD simulations. | S2 |
| **Figure S2:** Frequency of occurrence of eIF4E interactions with 4EGI-1 and i4EG-BiP in the frontal binding mode. | S3 |
| **Figure S3:** Backbone RMSD of in silico designed eIF4E mutants. | S4 |
| **Figure S4:** Expression and cap-binding activity of wild-type and mutant eIF4E proteins. | S5 |
| **Figure S5:** Determination of binding affinities of 4EGI-1 and i4EG-BiP for eIF4E mutants. | S6 |
| **Figure S6:** Protein production, cap-binding assay, and $K_D$ measurements for eIF4E double mutants with 4EGI-1 and i4EG-BiP. | S7 |
| **Figure S7**: Convergence of Funnel Metadynamics calculations on 4EGI-1. | S8 |
| **Figure S8:** Convergence of Funnel Metadynamics calculations on i4EG-BiP. | S9 |
| **Figure S9:** Alignment of the wild-type sequences of the eIF4G and 4E-BP1 peptides. | S10 |
| **Figure S10:** Comparative contacts analysis between the non-canonical motif of eIF4G and 4E-BP1 with apo eIF4E. | S11 |
| **Figure S11:** RMSD analysis of eIF4E in complex with eIF4G and 4E-BP1 peptides. | S12 |
| **Figure S12:** Residues of eIF4E and 4E-BP1 showing interaction changes upon ligand binding. | S13 |
| **Figure S13:** RMSD of 4EGI-1 and i4EG-BiP during MD simulations in complex with eIF4E, with or without 4E-BP1. | S14 |
| **Figure S14:** Torsional energy scan of the ad hoc parameterized dihedral angle in 4EGI-1 and i4EG-BiP. | S15 |
| **Figure S15:** RMSD analysis of $m^7$GTP binding across all simulations. | S16 |
| **Figure S16**: Tridimensional representation of the funnel-shaped restraining potential used in Funnel Metadynamics calculations. | S17 |
| **Table S1:** Residue conservation score of the eIF4E binding sites. | S18 |
| **Table S2:** Frequency of occurrence of contacts between eIF4E and the canonical and linker motifs of the 4E-BP1 peptide. | S19 |
| **Table S3:** Frequency of occurrence of contacts between eIF4E and the canonical and linker motifs of the eIF4G peptide. | S20 |
| **Table S4:** Summary of the MD simulations performed. | S21 |

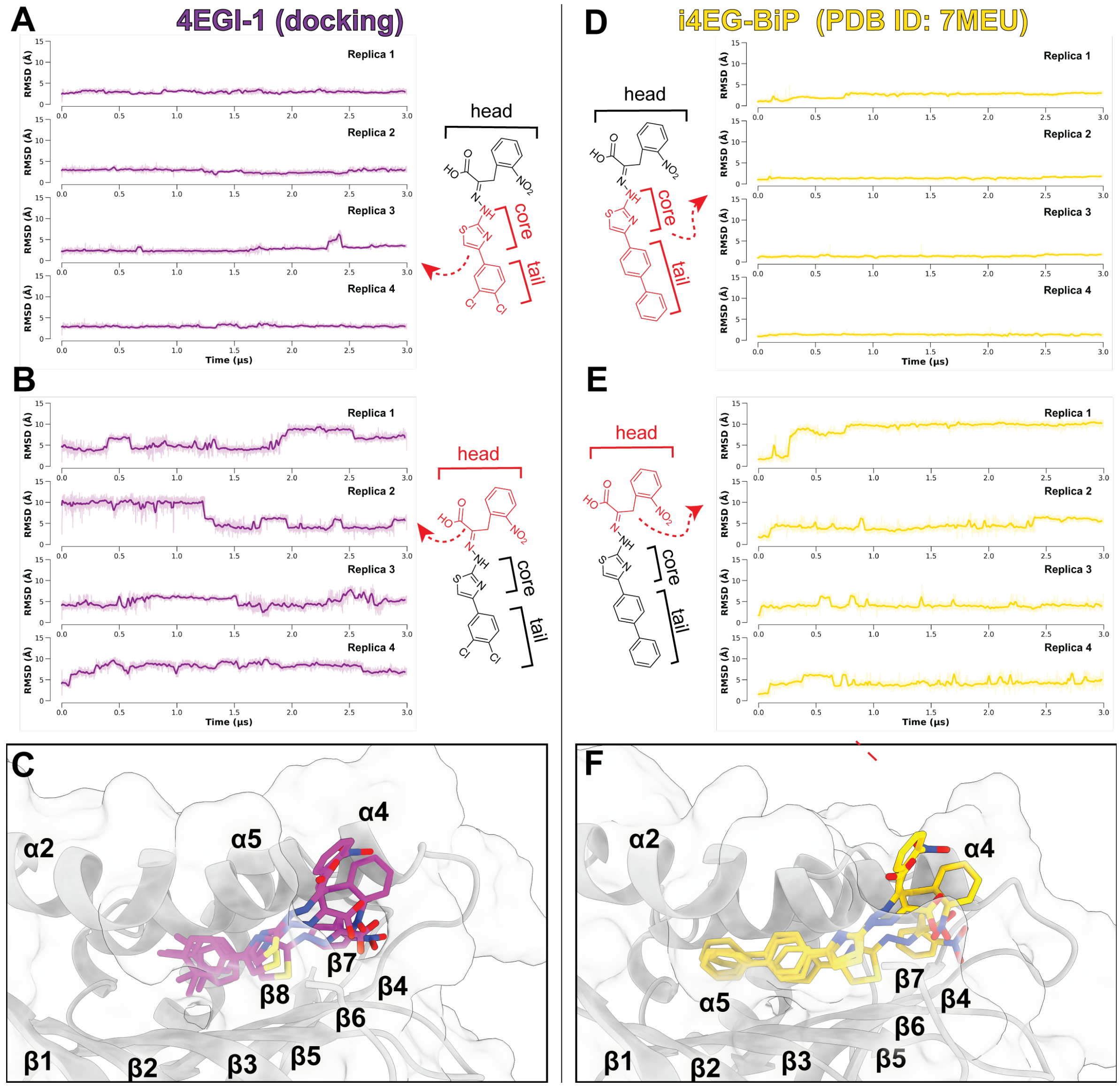

**Figure S1.** *Conformational dynamics of the frontal binding mode of 4EGI-1 and i4EG-BiP during MD simulations.* (A, D) RMSD of core plus tail heavy atoms for 4EGI-1 (A, magenta lines) and i4EG-BiP (D, yellow lines), computed over four independent 3 µs simulations. (B, E) RMSD of head group heavy atoms for 4EGI-1 (B, magenta lines) and i4EG-BiP (E, yellow lines), computed as above. 5 ns rolling averages are shown as bold lines, while raw data are shown as faded lines. Analyzed atoms are highlighted in red in the corresponding 2D ligand representation. (C, F) The three most representative poses (accounting for ~70% of the simulation frames) of (C) 4EGI-1 (magenta sticks) and (F) i4EG-BiP (yellow sticks) within the frontal binding site (PDB ID: 7MEU[1]), obtained by clustering the combined 12 µs MD trajectories using an RMSD cutoff of 1.5 Å. eIF4E is shown as gray transparent cartoon and surface.

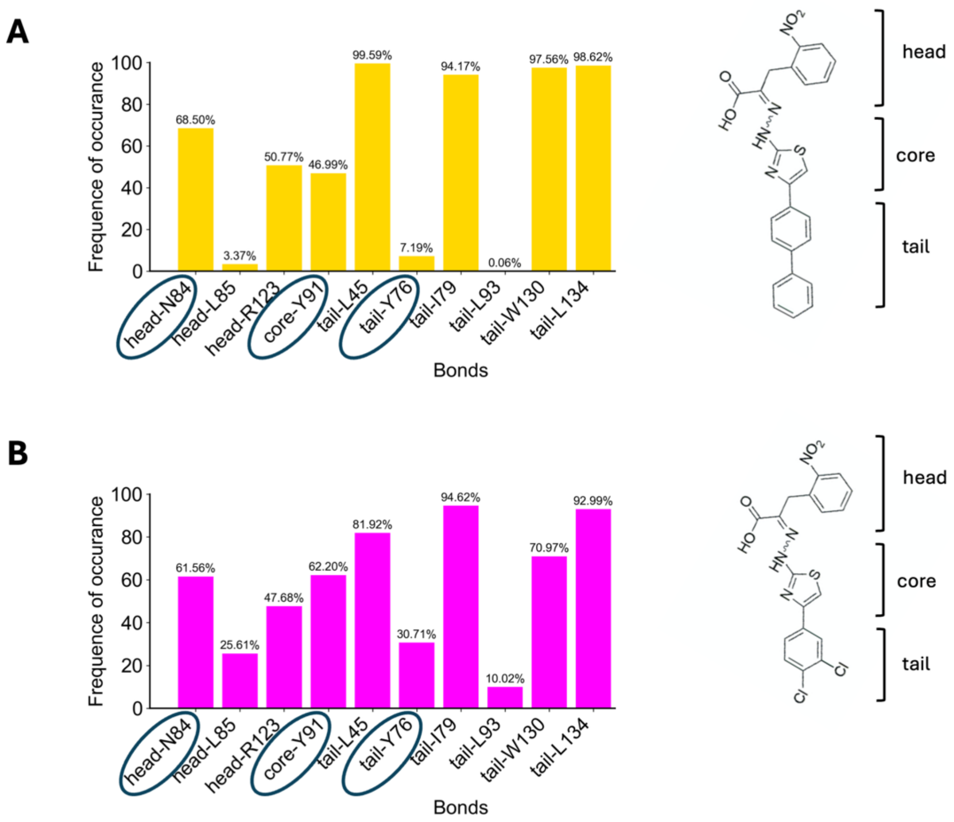

**Figure S2.** *Frequency of occurrence of eIF4E interactions with 4EGI-1 and i4EG-BiP in the frontal binding mode.* Bar plots report the frequency of occurrence (%) of key ligand–protein contacts observed over 12 µs of MD simulations for the eIF4E/4EGI-1 (A) and eIF4E/i4EG-BiP (B) complexes. Interacting residues are grouped by ligand regions (head, core, tail), as illustrated in the adjacent 2D structures. A 5.0 Å cutoff was applied for N84 and W130, and 5.5 Å for all other residues.

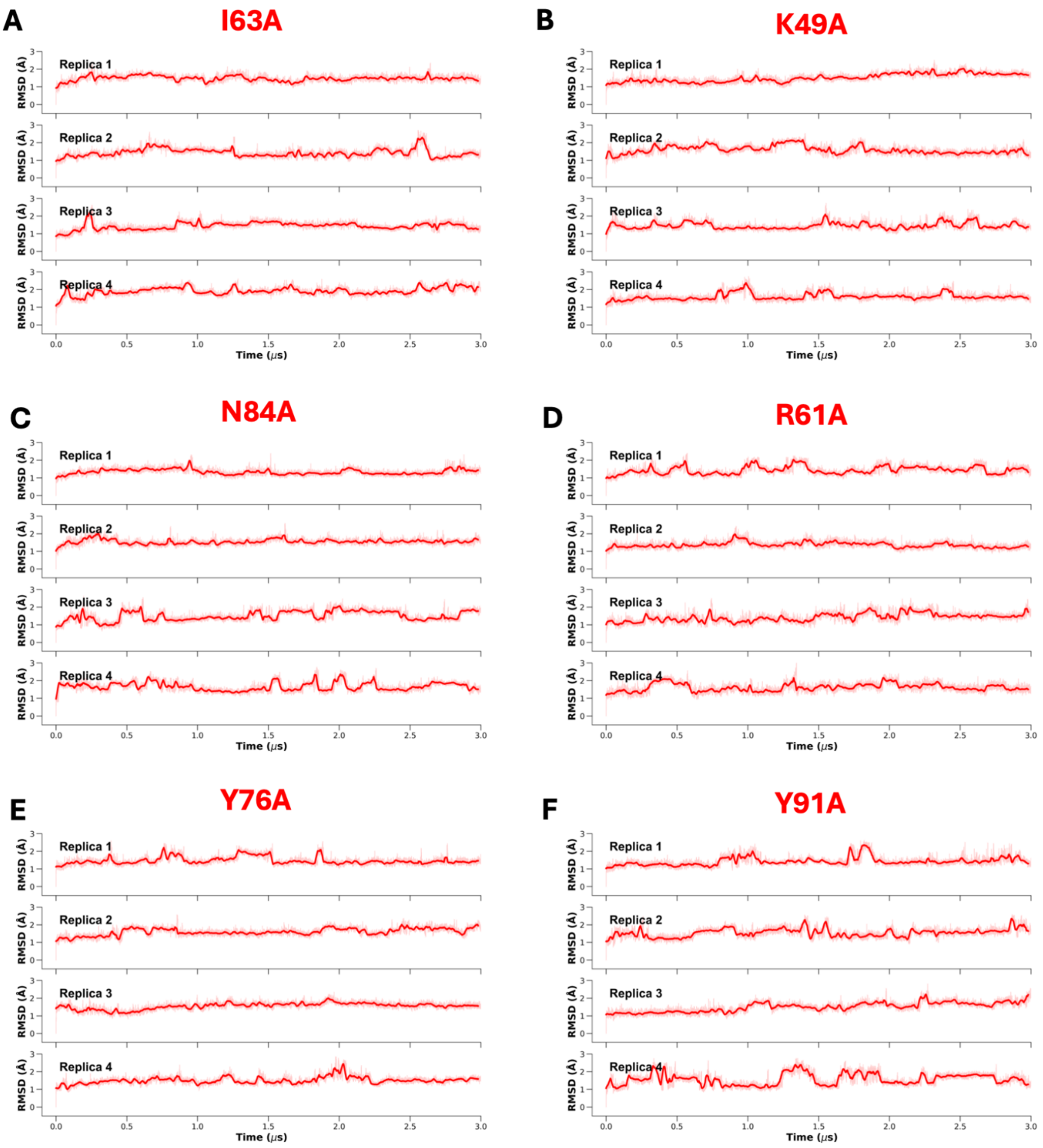

**Figure S3.** *Backbone RMSD of in silico designed eIF4E mutants.* Backbone RMSD of the six single-point eIF4E mutants (I63A, K49A, N84A, R61A, Y76A, Y91A) in their apo form, over four independent 3 μs MD simulations. Data are shown as a 5 ns rolling average (bold lines) and raw data (faded lines).

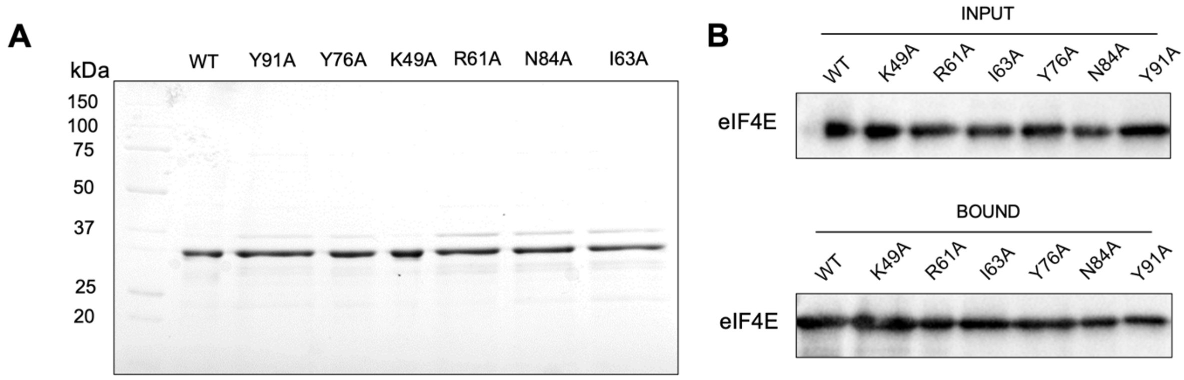

**Figure S4.** *Expression and cap-binding activity of wild-type and mutant eIF4E proteins.* (A) SDS-PAGE analysis on a 12.5% polyacrylamide gel of 2 μg of purified eIF4E WT and single-point mutants K49A, R61A, I63A, Y76A, N84A and Y91A. (B) Cap-binding activity was assessed via $m^7$GTP pull-down assay followed by Western blot analysis: the upper panel shows the input protein (0.5 μg), while the lower panel displays the bound fractions.

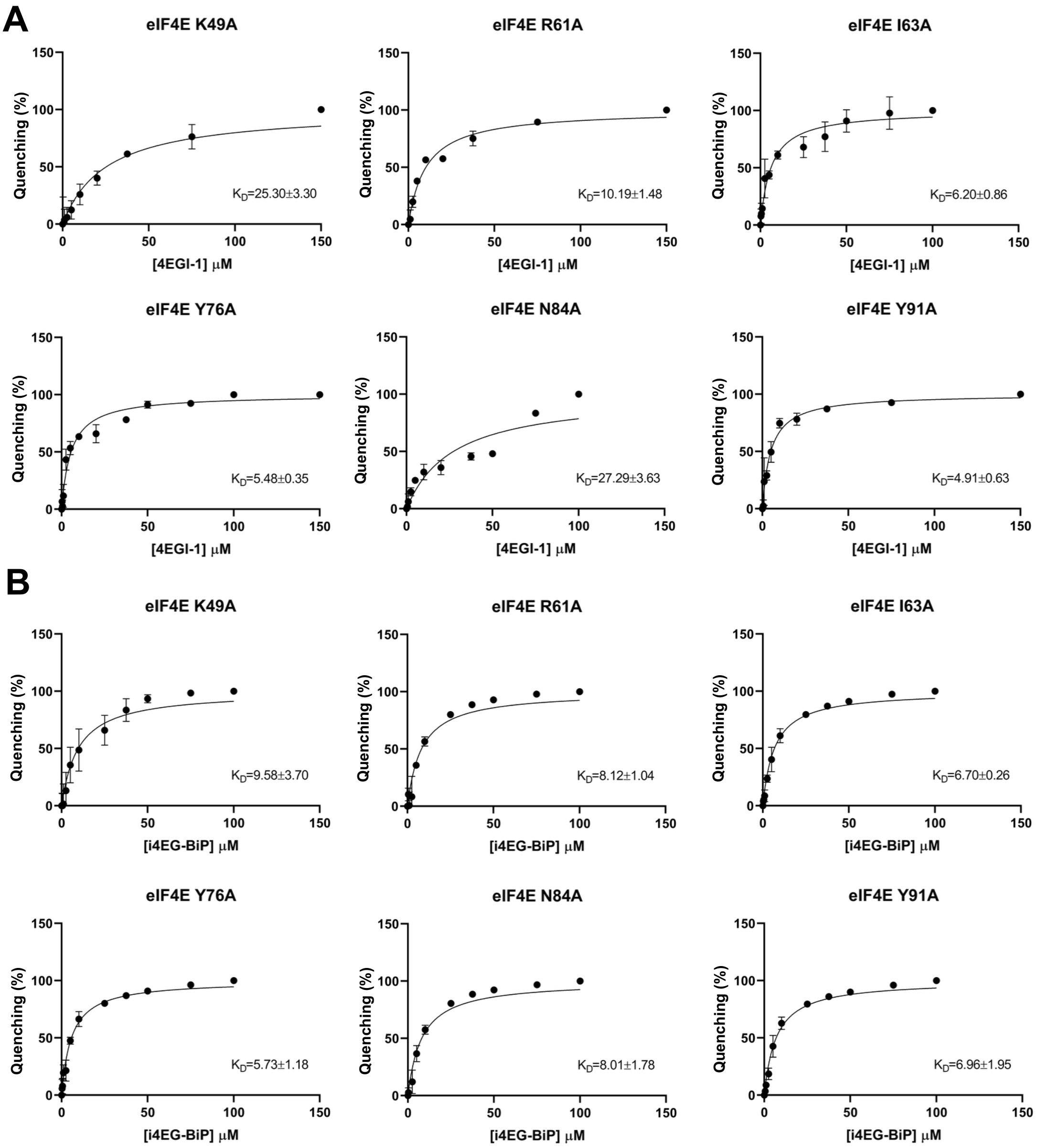

**Figure S5.** *Determination of binding affinities of 4EGI-1 and i4EG-BiP for eIF4E mutants.* The plots show data from fluorescence quenching assays on eIF4E mutants with increasing concentrations of either 4EGI-1 (A) or i4EG-BiP (B). Data are expressed as percentage of intrinsic fluorescence intensity quenching versus inhibitor concentration and were fitted using a one-site specific binding model. Mean $K_D$ ± standard deviation (SD) values are reported from three independent experiments; error bars indicate SD.

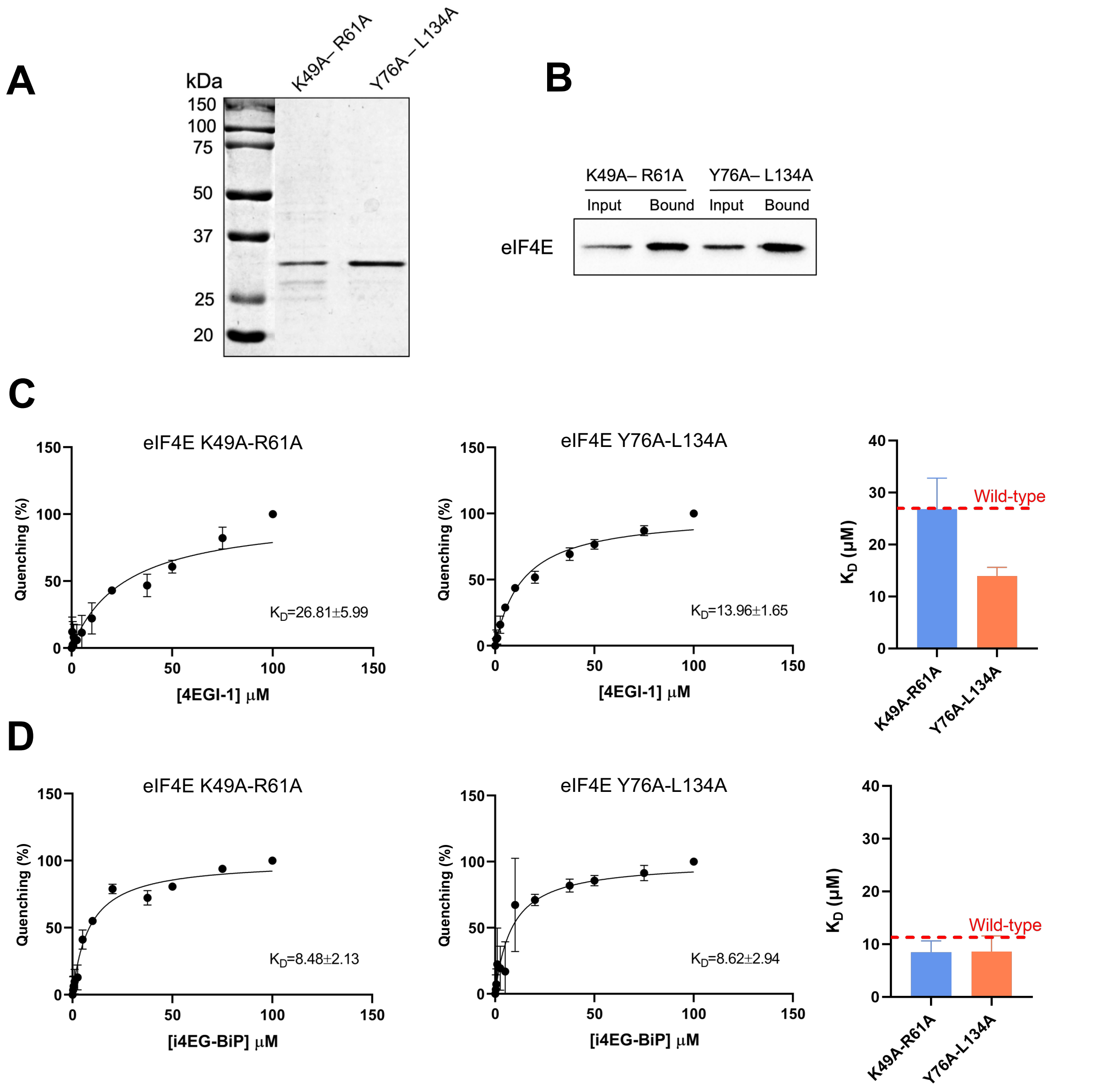

**Figure S6.** *Protein production, cap-binding assay, and $K_D$ measurements for eIF4E double mutants with 4EGI-1 and i4EG-BiP.* (A) 2 µg of purified eIF4E double mutants (K49A-LR61A and Y76A-L134A) were separated on a 12.5% SDS-PAGE gel. (B) Cap-binding activity was evaluated by $m^7$GTP affinity pull-down followed by Western blot analysis; input (0.5 µg) and bound fractions are shown. (C, D) Fluorescence quenching assays performed by titrating eIF4E double-mutants with increasing concentrations of either 4EGI-1 (C) or i4EG-BiP (D). Data are shown as percentage of intrinsic fluorescence intensity quenching versus inhibitor concentration and fitted using a one-site specific binding model. Mean $K_D$ ± SD values are from three independent experiments; error bars represent SD. Bar plots summarize $K_D$ values of eIF4E double mutants with the $K_D$ of WT eIF4E shown as a reference by a red dotted line.

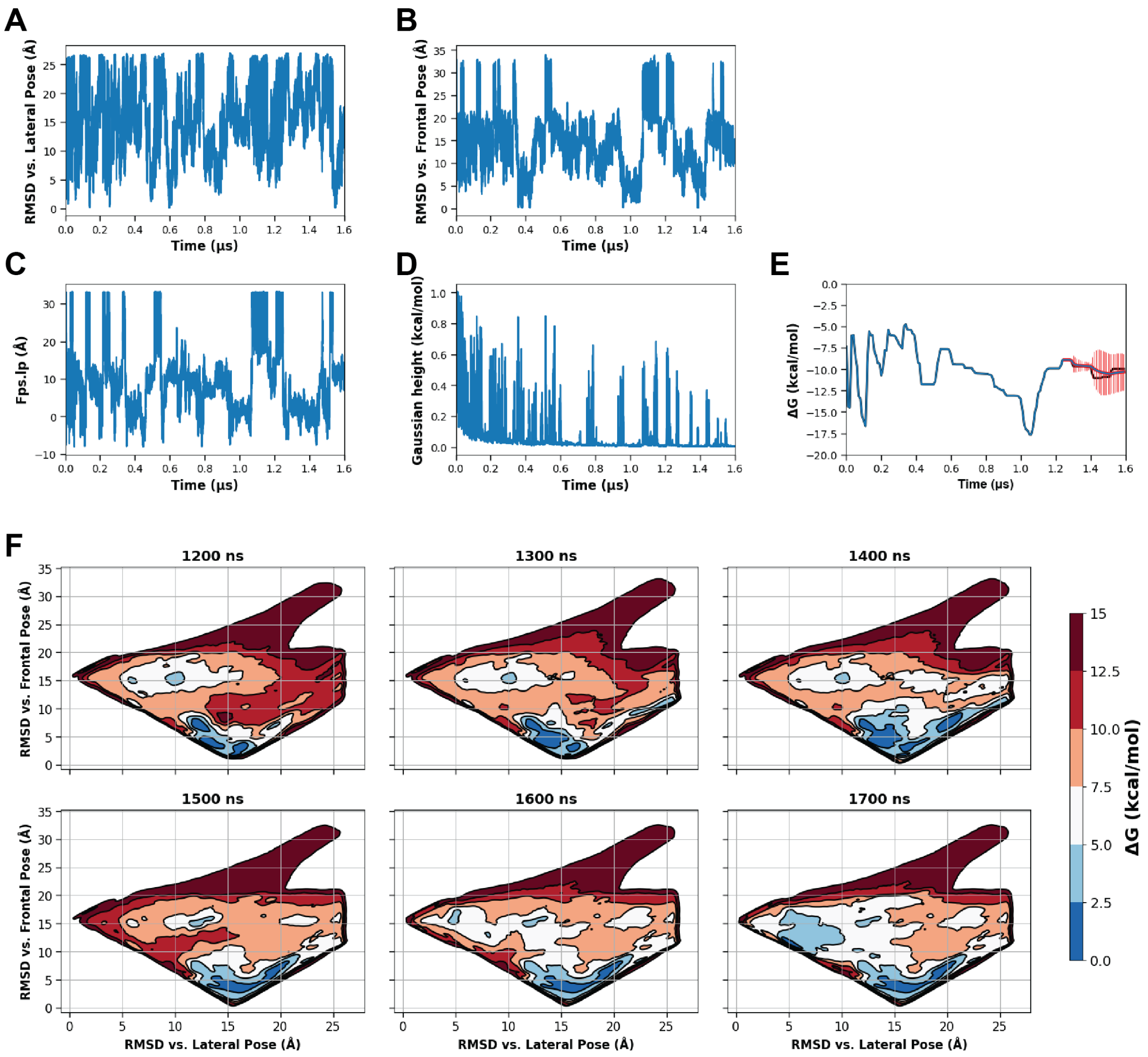

**Figure S7.** *Convergence of Funnel Metadynamics calculations on 4EGI-1.* (A–C) Time evolution of the key collective variables (CVs): (A) RMSD of 4EGI-1 relative to the lateral binding pose, (B) RMSD relative to the frontal binding pose, and (C) projection of the ligand's center of mass along the funnel axis (*fps.lp*). (D) Gaussian hills heights over simulation time, illustrating the progressive decrease in bias deposition typical of the well-tempered metadynamics. (E) Estimated binding free energy ($\Delta G$) as a function of time, with a time-weighted mean (red line) and SD (shaded area) calculated over 100 ns sliding windows in the 1.3-1.7 μs interval. (F) Free energy surface (FES) snapshots taken every 100 ns throughout the convergence phase, projected on RMSD from the lateral (x-axis) and frontal (y-axis) poses. The stabilization of the main energy basin supports convergence toward the frontal binding mode.

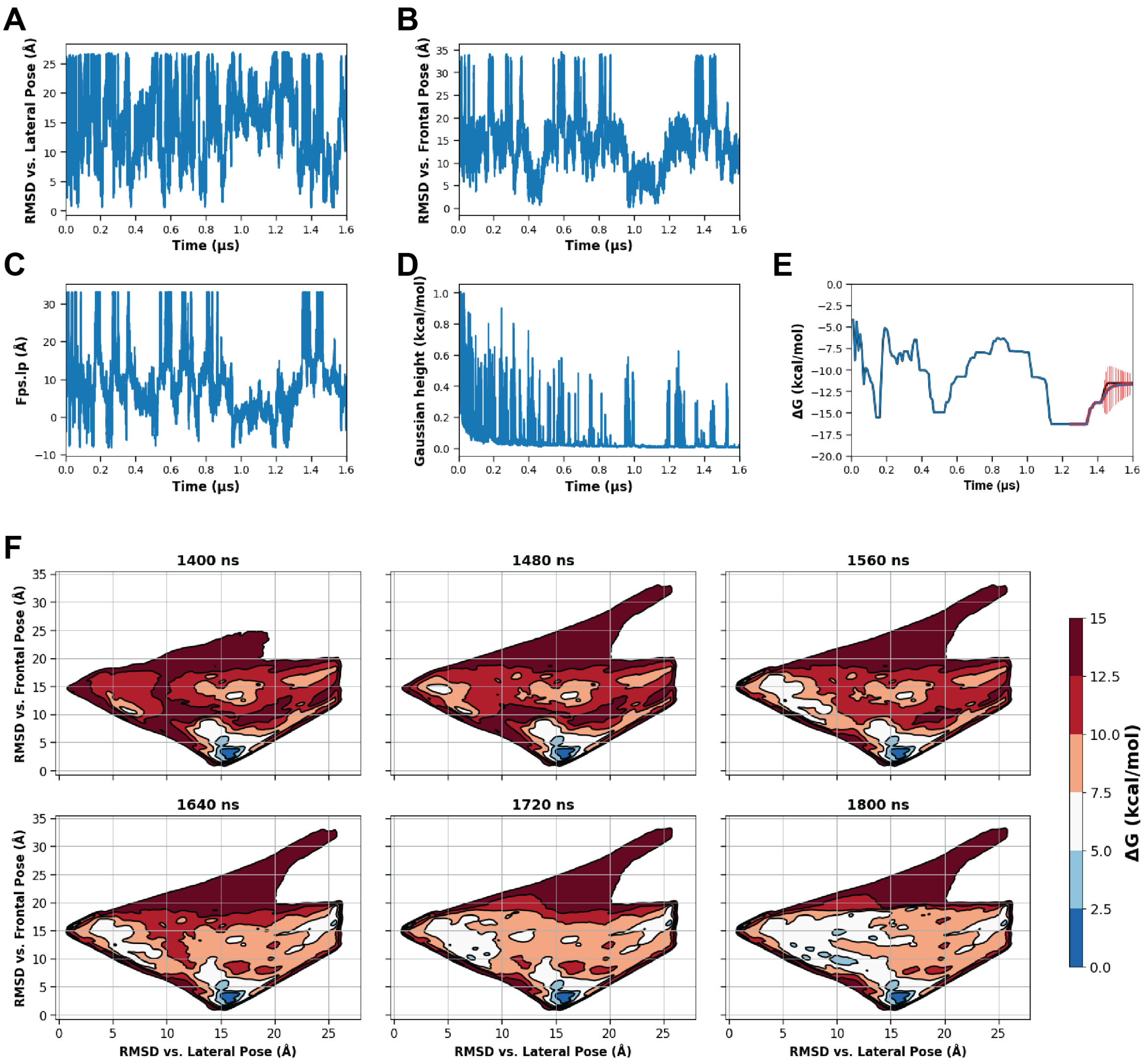

**Figure S8.** *Convergence of Funnel Metadynamics calculations on i4EG-BiP.* (A–C) Time evolution of the key collective variables (CVs): (A) RMSD of 4EGI-1 relative to the lateral binding pose, (B) RMSD relative to the frontal binding pose, and (C) projection of the ligand's center of mass along the funnel axis (*fps.lp*). (D) Gaussian hills heights over simulation time, illustrating the progressive decrease in bias deposition typical of the well-tempered metadynamics. (E) Estimated binding free energy ($\Delta G$) as a function of time, with a time-weighted mean (red line) and SD (shaded area) calculated over 100 ns sliding windows in the 1.4-1.8 µs interval. (F) Free energy surface (FES) snapshots taken every 100 ns throughout the convergence phase, projected on RMSD from the lateral (x-axis) and frontal (y-axis) poses. The stabilization of the main energy basin supports convergence toward the frontal binding mode.

```
                  CM           Linker        NCM
4E-BP1 50   TRIIYDRKFLMECRNSPVTKTPPRDLPTIPGVTSP 84
            .:..|||:||:..:....:...|..||.|..|...
eIF4G  608  EKKRYDREFLLGFQFIFASMQKPEGLPHISDVVLD 642
```

**Figure S9.** *Alignment of the wild-type sequences of the eIF4G and 4E-BP1 peptides.* Sequence alignment between the 4E-BP1 (PDB ID: 5BXV[2,3]) and eIF4G (PDB ID: 5T46) peptides was performed using the Needleman-Wunsch algorithm, using EMBL EMBOSS Needle tool. According to the software, markup line uses ":" for a similarity which scores more than 1.0, "|" for an identity where both sequences have the same residue and "." for positive scores that are minor than 1.0 and "-" for gaps (not present here). Amino acids are grouped into the Canonical Motif (CM), Linker, and Non-Canonical Motif (NCM). The conserved YXXXXLΦ motif typical of 4E binding proteins is highlighted in yellow.

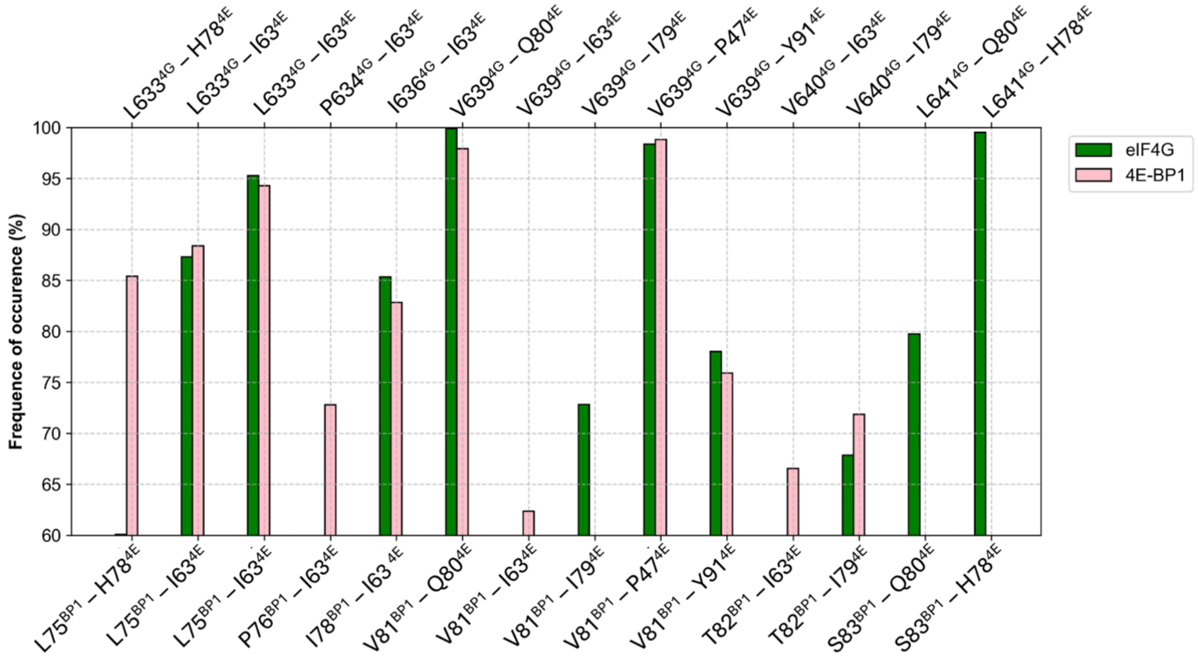

**Figure S10.** *Comparative contacts analysis between the non-canonical motif of eIF4G and 4E-BP1 with apo eIF4E.* Bar plot showing the frequency of occurrence of contacts between eIF4E and the non-canonical motifs (NCM) of the eIF4G (green bars; labels above the graph), or the 4E-BP1 peptides (pink bars; labels below the graph). Only contacts with a frequency of occurrence greater than 60% are reported.

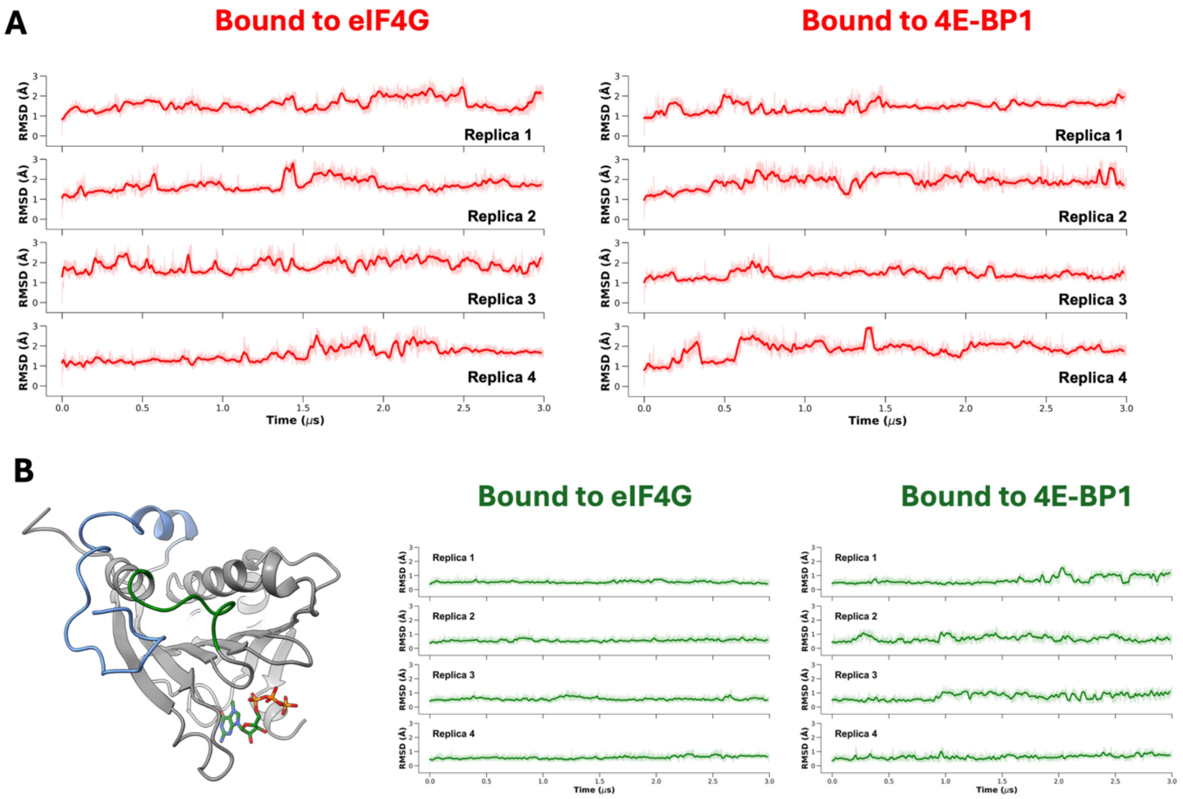

**Figure S11.** *RMSD analysis of eIF4E in complex with eIF4G and 4E-BP1 peptides.* (A) Backbone RMSD of eIF4E over four independent 3 μs MD simulations in complex with eIF4G (left panel, PDB ID: 5T46[2]) and 4E-BP1 (right panel, PDB ID: 5BXV[2]). RMSD traces are shown as 5 ns rolling average (bold lines) and raw data (faded lines). (B) Backbone RMSD of the eIF4E segment encompassing residues 77 to 85 across the same replicas in complex with eIF4G (left panel) and 4E-BP1 (right panel), shown as 5 ns rolling average (bold lines) and raw data (faded lines). In the 3D protein representation, the amino acids selected for the calculation are shown as green cartoon, while the reference 4E-BP1 peptide is depicted in light blue; $m^7GTP$ is shown as green sticks.

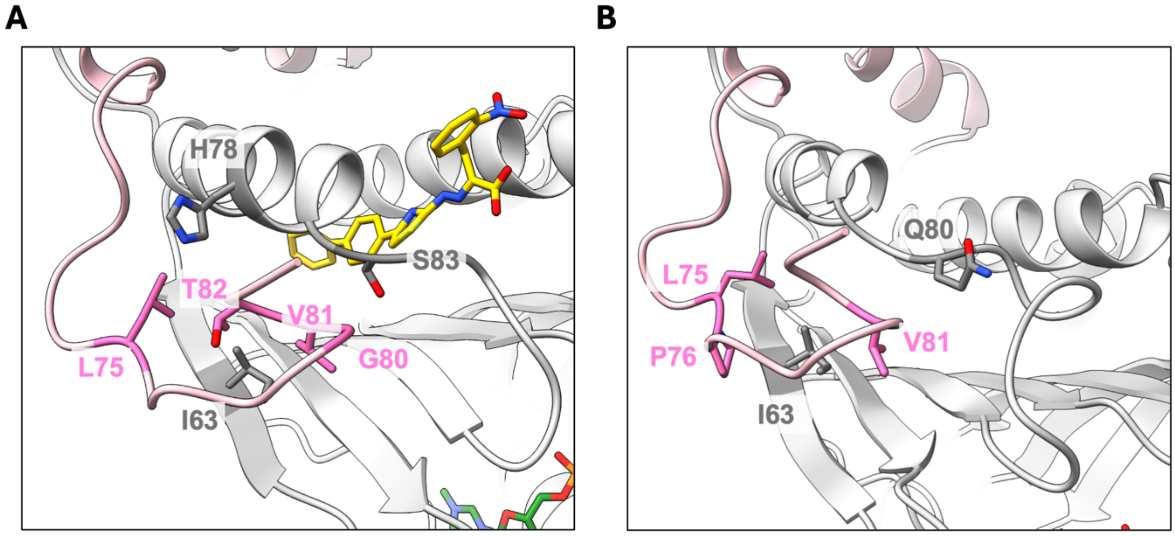

**Figure S12.** *Residues of eIF4E and 4E-BP1 showing interaction changes upon ligand binding.* (A) Residues involved in new interactions (G80$^{BP1}$-S83$^{4E}$, T82$^{BP1}$-H78$^{4E}$) and in contacts with increased frequency (L75 $^{BP1}$-H78$^{4E}$, V81 $^{BP1}$-I63$^{4E}$, T82 $^{BP1}$-I63$^{4E}$) in the ligand-bound eIF4E/4E-BP1 complex. (B) Residues involved in native interactions that are lost (V81$^{BP1}$-Q80$^{4E}$) or reduced in frequency (L75$^{BP1}$-I63$^{4E}$, P76$^{BP1}$-I63$^{4E}$) compared to the apo complex. As a representative example, only the structure of the eIF4E/4E-BP1 complex bound to i4EG-BiP is shown. The 4E-BP1 peptide is shown in pink, eIF4E in grey, i4EG-BiP in yellow, and m$^7$GTP in green. Proteins are displayed as ribbons; interacting residues and i4EG-BiP are highlighted in darker shades and shown as sticks. The two representative structures correspond to the centroid of the dominant cluster (1.5 Å RMSD cutoff) from 12 µs of simulation.

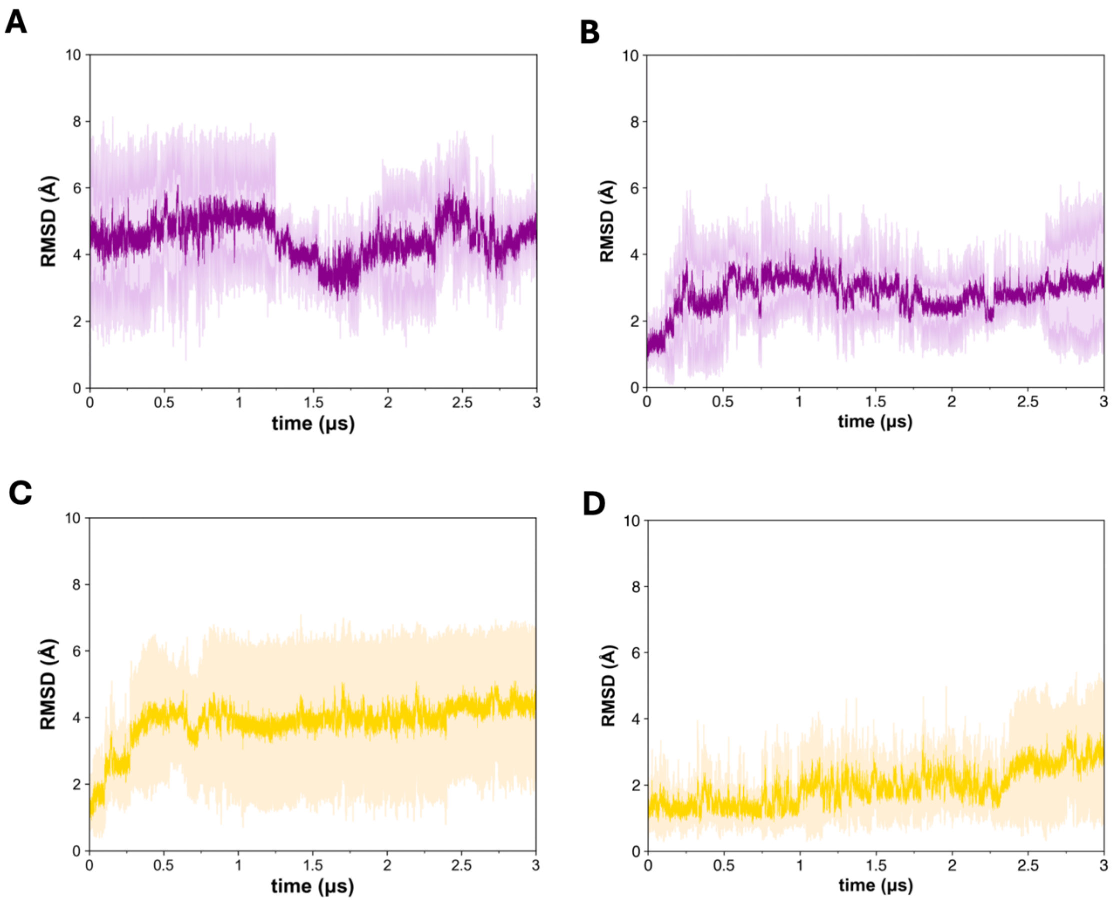

**Figure S13.** *RMSD of 4EGI-1 and i4EG-BiP during MD simulations in complex with eIF4E, with or without 4E-BP1.* (A–B) RMSD of heavy atoms of 4EGI-1 in the frontal binding pose, in complex with eIF4E alone (A) or with eIF4E bound to 4E-BP1 (B). (C–D) RMSD of heavy atoms of i4EG-BiP in the frontal binding pose, in complex with eIF4E alone (C) or with eIF4E bound to 4E-BP1 (D). Data are shown as raw values (faded lines) and 5 ns rolling averages (bold lines).

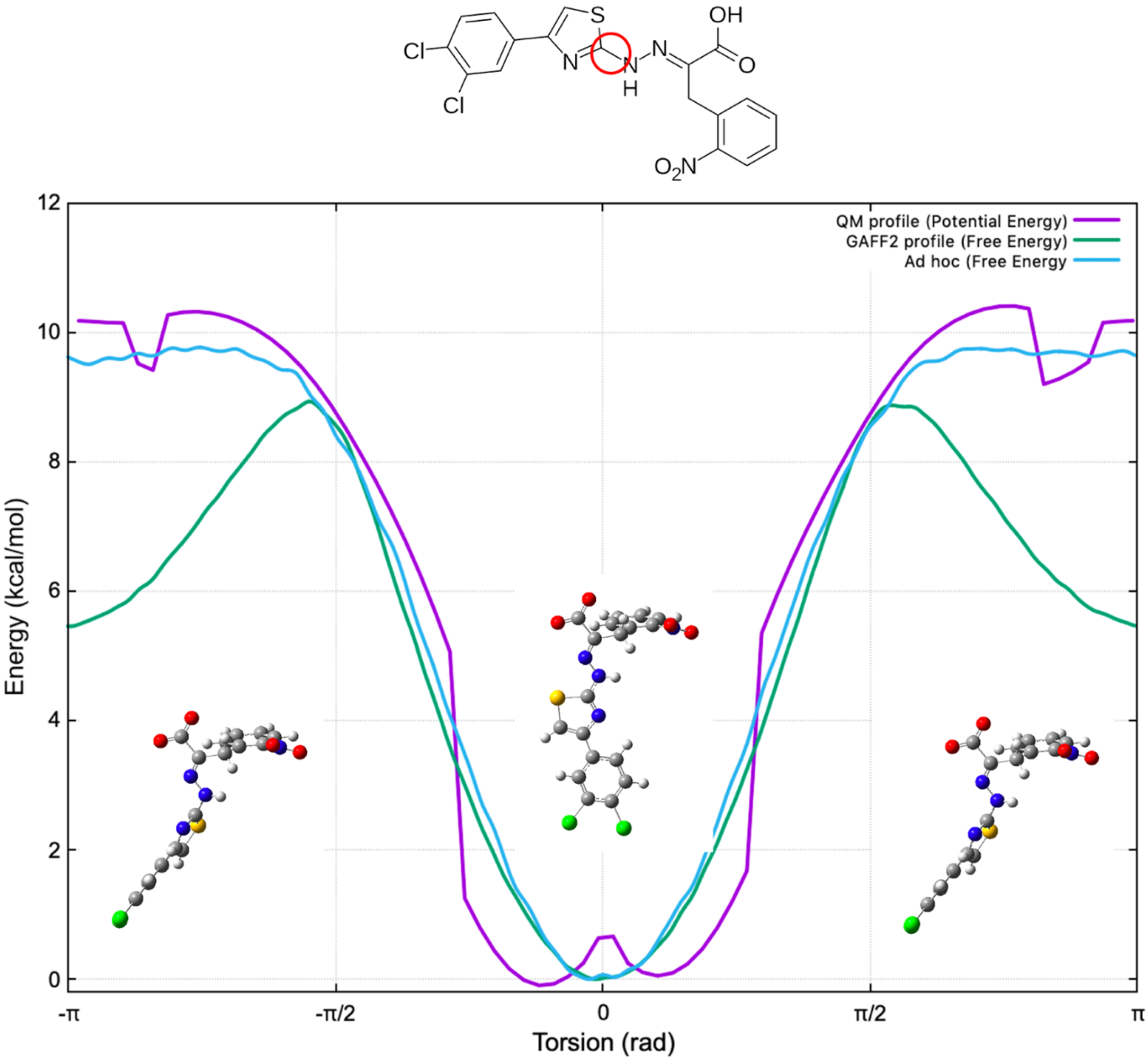

**Figure S14.** *Torsional energy scan of the ad hoc parameterized dihedral angle in 4EGI-1 and i4EG-BiP.* Comparison of torsional energy profiles obtained from quantum mechanical (QM) calculations (magenta), the original GAFF2 force field parameters (cyan), and the optimized ad hoc parameters (green) for the dihedral highlighted in red. The plot shows the potential or free energy as a function of the torsional angle (in radians), illustrating the improved agreement between the ad hoc parametrization and the QM reference. Molecular conformations at selected torsion angles are shown below the graph.

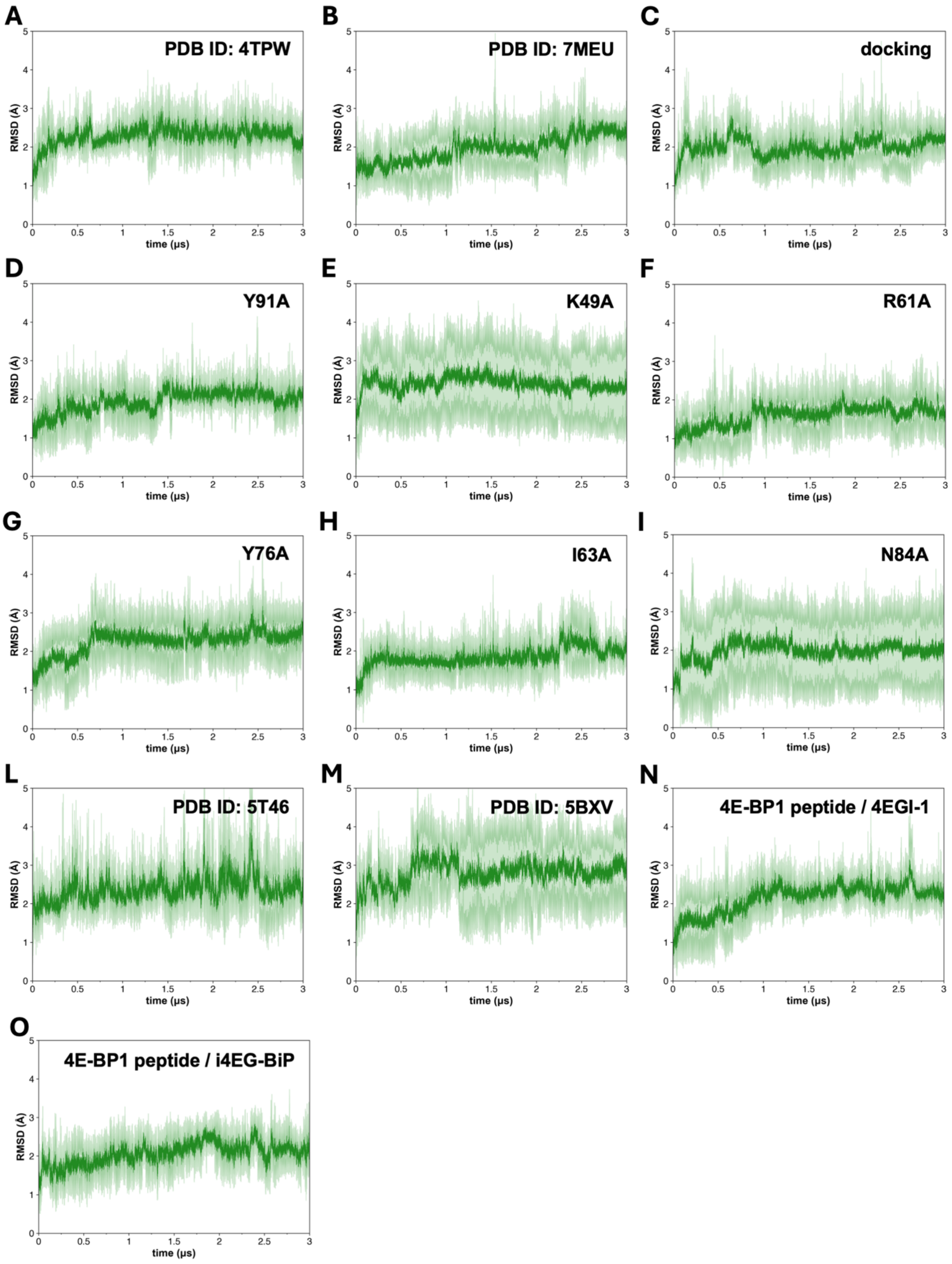

**Figure S15.** *RMSD analysis of m$^7$GTP binding across all simulations.* Heavy atoms RMSD of m$^7$GTP averaged over four independent 3 μs simulations for each analyzed system: (A) eIF4E/4EGI-1 complex (PDB ID: 4TPW[4]), (B) eIF4E/i4EG-BiP complex (PDB ID: 7MEU[1]), (C) eIF4E in complex with 4EGI-1 docked at the frontal site, (D-I) eIF4E single-mutants, (L) eIF4E/eI4EG complex(PDB ID: 5T46)[3], (M) eIF4E/4E-BP1 complex (PDB ID: 5BXV[2]), (N) eIF4E/4E-BP1/4EGI-1 ternary complex, (O) eIF4E/4E-BP1/i4EG-BiP ternary complex. RMSD traces are shown as rolling averages (dark lines) with shaded areas representing standard deviation.

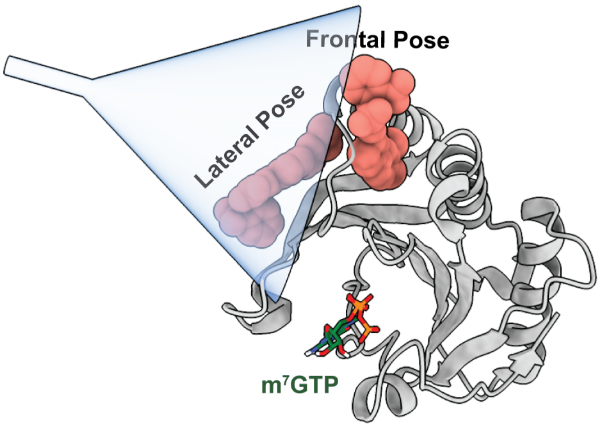

**Figure S16.** Tridimensional representation of the funnel-shaped restraining potential used in Funnel Metadynamics calculations.

**Table S1. Residue conservation score of the eIF4E binding residues.** Conservation scores of residues located at the frontal and lateral sites of eIF4E, calculated using ConSurf webserver (consurf.tau.ac.il). Scores range from 1 (variable) to 9 (conserved).

| Residue | Conservation index | Binding site |
|---|---|---|
| L45 | 6 | frontal |
| Y76 | 6 | frontal |
| I79 | 8 | frontal |
| N84 | 3 | frontal |
| L85 | 8 | frontal |
| Y91 | 8 | frontal |
| L93 | 6 | frontal |
| R123 | 1 | frontal |
| **W130** | **9** | **frontal** |
| L134 | 5 | frontal |
| K49 | 6 | lateral |
| R61 | 5 | lateral |

**Table S2. Frequency of occurrence of contacts between eIF4E and the canonical and linker motifs of the 4E-BP1 peptide.** The table reports the frequency of occurrence (%) of individual residue contacts observed between eIF4E and the canonical motif (CM) or the linker region of the 4E-BP1 peptide during MD simulations. The CM interacts with the dorsal surface of eIF4E, as seen in the crystal structure of the complex (PDB ID: 5BXV), involving key contacts such as R63, Y54, and R56 of 4E-BP1 with H47, L39, V69, and W73 of eIF4E. Linker interactions are primarily mediated by T68, V67, and P71 with eIF4E residues N77 and H78[2,5].

Additional native contacts not previously described in the literature but consistently observed in our simulations are also included.

| 4E-BP1 Motif | Contact | Frequence of occurrence (%) |
|---|---|---|
| Canonical | $R63^{BP1}$-$W73^{4E}$ | 99.98 |
| Canonical | $L59^{BP1}$-$W73^{4E}$ | 99.91 |
| Canonical | $R56^{BP1}$-$E132^{4E}$ | 99.87 |
| Canonical | $L59^{BP1}$-$V69^{4E}$ | 99.86 |
| Canonical | $C62^{BP1}$-$V69^{4E}$ | 99.85 |
| Canonical | $I52^{BP1}$-$Q40^{4E}$ | 99.66 |
| Canonical | $L59^{BP1}$-$L135^{4E}$ | 98.53 |
| Canonical | $R51^{BP1}$-$D147^{4E}$ | 96.33 |
| Canonical | $F58^{BP1}$-$H47^{4E}$ | 95.93 |
| Canonical | $Y54^{BP1}$-$V69^{4E}$ | 94.52 |
| Canonical | $M60^{BP1}$-$W73^{4E}$ | 93.88 |
| Canonical | $R51^{BP1}$-$Q40^{4E}$ | 91.17 |
| Canonical | $L59^{BP1}$-$I138^{4E}$ | 90.53 |
| Canonical | $M60^{BP1}$-$L135^{4E}$ | 85.67 |
| Canonical | $I53^{BP1}$-$Q140^{4E}$ | 85.42 |
| Canonical | $Y54^{BP1}$-$G139^{4E}$ | 85.33 |
| Canonical | $Y54^{BP1}$-$L39^{4E}$ | 84.97 |
| Canonical | $R51^{BP1}$-$Q140^{4E}$ | 83.84 |
| Canonical | $C62^{BP1}$-$W73^{4E}$ | 81.54 |
| Linker | $T68^{BP1}$-$W73^{4E}$ | 99.66 |
| Linker | $V67^{BP1}$-$E70^{4E}$ | 98.97 |
| Linker | $P71^{BP1}$-$H78^{4E}$ | 93.78 |
| Linker | $T68^{BP1}$-$N77^{4E}$ | 86.26 |
| Linker | $P72^{BP1}$-$L75^{4E}$ | 83.14 |
| Linker | $T68^{BP1}$-$E70^{4E}$ | 76.05 |

**Table S3. Frequency of occurrence of contacts between eIF4E and the canonical and linker motifs of the eIF4G peptide.** The table reports the frequency of occurrence (%) of residue contacts observed between eIF4E and the canonical motif (CM) or the linker region of the eIF4G peptide during MD simulations. The CM binds the dorsal surface of eIF4E, as shown in the crystal structure of the complex (PDB ID: 5T46), with Y612, L617, and L618 of eIF4G forming key contacts with eIF4E residues H37, P38, L39, V69, W73, L135, and E132. In the linker region, Q621, S626, L641, and K629 interact mainly with H78 and N77 of eIF4E[3]. Additional native contacts not previously described in the literature but consistently observed in our simulations are also included.

| eIF4G Motif | Contact | Frequence of occurrence (%) |
|---|---|---|
| Canonical | $R614^{4G}$ -$E132^{4E}$ | 100 |
| Canonical | $Y612^{4G}$ -$P38^{4E}$ | 99.92 |
| Canonical | $L617^{4G}$ -$V69^{4E}$ | 99.83 |
| Canonical | $L617^{4G}$ -$W73^{4E}$ | 99.76 |
| Canonical | $K610^{4G}$ -$Q40^{4E}$ | 99.5 |
| Canonical | $P620^{4G}$ -$H37^{4E}$ | 99.23 |
| Canonical | $P620^{4G}$ -$V69^{4E}$ | 99.18 |
| Canonical | $L617^{4G}$ -$L135^{4E}$ | 98.84 |
| Canonical | $P620^{4G}$ -$W73^{4E}$ | 98.8 |
| Canonical | $L618^{4G}$ -$W73^{4E}$ | 96.46 |
| Canonical | $L617^{4G}$ -$I138^{4E}$ | 94.98 |
| Canonical | $Y612^{4G}$ -$G139^{4E}$ | 90.46 |
| Canonical | $L618^{4G}$ -$L135^{4E}$ | 88.11 |
| Canonical | $R611^{4G}$ -$E140^{4E}$ | 87.67 |
| Canonical | $Y612^{4G}$-$L39^{4E}$ | 84.63 |
| Linker | $K629^{4G}$-$H78^{4E}$ | 95.99 |
| Linker | $Q621^{4G}$-$N77^{4E}$ | 95.52 |
| Linker | $S626^{4G}$-$N77^{4E}$ | 94.04 |
| Linker | $P630^{4G}$-$A74^{4E}$ | 91.02 |

**Table S4. Summary of the MD simulations performed.** Overview of the simulated systems, including details on the protein constructs, binding partners, initial structural models (e.g., X-ray or docked conformations), and total simulation times for each condition.

| Binding partners | Starting structure | Simulation time |
|---|---|---|
| $m^7GTP$, 4EGI-1 | X-ray (PDB ID: 4TPW) | 3 μs x 4 replicas standard MD and 1.7 μs FM |
| $m^7GTP$, i4EG-BiP | X-ray (PDB ID: 7MEU) | 3 μs x 4 replicas standard MD and 1.8 μs FM |
| $m^7GTP$, 4EGI-1 | Docking of 4EGI-1 in the eIF4E/i4EG-BiP X-ray structure (PDB ID: 7MEU) | 3 μs x 4 replicas standard MD |
| $m^7GTP$ | X-ray (PDB ID: 1IPC) | 3 μs x 4 replicas standard MD |
| $m^7GTP$ | Modified X-ray (PDB ID: 1IPC) | 3 μs x 4 replicas standard MD x 6 mutants |
| $m^7GTP$, eIF4G peptide | X-ray (PDB ID: 5T46) | 3 μs x 4 replicas standard MD |
| $m^7GTP$, 4E-BP1 peptide | X-ray (PDB ID: 5BXV) | 3 μs x 4 replicas standard MD |
| $m^7GTP$, i4EG-BiP, 4E-BP1 peptide | 4E-BP1 MD pose assembled with the X-ray eIF4E/i4EG-BiP (PDB ID: 7MEU) | 3 μs x 4 replicas standard MD |
| $m^7GTP$, 4EGI-1, 4E-BP1 peptide | 4E-BP1 MD pose assembled with the eIF4E/4EGI-1 docking complex | 3 μs x 4 replicas standard MD |
